# *COCHLEATA* controls spatial regulation of cytokinin and auxin during nodule development

**DOI:** 10.1101/2025.08.22.671680

**Authors:** Karen Velandia, Muhammad Nouman Sohail, Tiana Scott, Alejandro Correa-Lozano, Alannah Mannix, Eloise Foo

**Affiliations:** Discipline of Biological Sciences, School of Natural Sciences, University of Tasmania, Private Bag 55, Hobart, Tasmania, 7001, Australia

**Keywords:** Autoregulation of nodulation, auxin, COCHLEATA, cytokinin, gibberellin, nodulation, root development

## Abstract

Root nodules develop in some legumes that host nitrogen-fixing bacteria and likely evolved through modifications of the ancestral lateral root program with plant hormones playing key regulatory roles. Members of the NOOT-BOP-COCH-LIKE transcriptional co-regulator family suppress root identity in legume nodules, including *Pisum sativum coch1* that display root-nodule hybrids. However, how COCH/NOOT interacts with hormones to control nodule organogenesis is unclear. We show that *PsCOCH* (*COCHLEATA*) is required for spatial tight regulation of auxin and cytokinin during nodule organogenesis and identify key hormone and signalling genes regulated by *COCH*. COCH suppresses cytokinin levels and response during nodule formation, as cytokinin levels are elevated in *Pscoch* abnormal nodules and this is mirrored by ectopic cytokinin-responsive *TCSn::GUS* expression in *Pscoch* nodule apices, nodule vasculature and in root-like tissue. In contrast, *PsCOCH* promotes auxin accumulation and precise auxin response patterning in nodules, as *Pscoch* mutants show significantly reduced auxin levels and severely altered auxin-responsive DR5::GUS expression patterns. RNAseq analysis revealed that *Pscoch* developing nodules have gene expression profiles more similar to root primordia, with increased expression of defence and auxin response genes (IAA and ARF) and reduced expression of cytokinin biosynthesis genes (*IPT3, CYP735A* and *LOG2)* compared to wild type. We found gibberellin is unlikely to act downstream of *PsCOCH*, as *Pscoch* and gibberellin-deficient double mutants still form root-nodule hybrids. Ectopic constitutive expression of *PsCOCH* also produces root-nodule hybrids and we found intriguing links between autoregulation of nodulation pathway and *COCH*, suggesting that a complex feedback mechanism acts in *COCH* control of nodule identity.

## Introduction

The formation of root nodules that host nitrogen-fixing bacteria in selected plants of the Fabid clade offer a unique advantage to these plants in nitrogen-poor environments and are also a significant nitrogen input into natural and agricultural environments. Decades of research in legume-rhizobial symbioses has uncovered many of the genes and hormonal signals controlling the two stages of legume nodulation, infection in the epidermis and nodule organogenesis in the inner root (Figure S3; Luo et al., 2023; Roy et al., 2020; Velandia et al., 2022). The infection program shares evolutionary history with the more ancient and widespread arbuscular mycorrhizal symbiosis (Radhakrishnan et al., 2020), while nodule organogenesis has overlaps with lateral root program including the cell layers that these organs arise from, as well as overlapping transcriptional networks and hormone dynamics (Lee et al., 2024; Schiessl et al., 2019; Soyano et al., 2019).

Intriguing players in the cross over of the root and nodule programs are the orthologous members of the NOOT-BOP-COCH-LIKE (NBCL) gene family *COCHLEATA/COCH2* in *Pisum sativum L.* and *NOOT/NOOT2* in *Medicago truncatula*. Pea *coch* and *Medicago noot* mutants display root-nodule hybrids (Couzigou et al., 2012; Ferguson and Reid, 2005), and act somewhat redundantly with *coch2* and *noot2* respectively to maintain nodule identity (Liu et al., 2023). The determinate nodulator *Lotus japonicus* only contains one NBCL member, *NBCL1*, and at least some *Ljnbc1* mutants display root-nodule hybrids, suggesting a conserved role for this gene in nodule organogenesis (Liu et al., 2018; Magne et al., 2018a). Studies in *Medicago* have revealed that *NOOT* likely functions by establishing boundaries between different cell populations within nodules to ensure proper cell fate specification (Shen et al., 2020b). This boundary formation is crucial for separating the infected tissue, where nitrogen fixation occurs, from the root vasculature. In addition, in *noot* mutants, there is a shift in the gradient of mitotic activity; pericycle and endodermis-derived cells exhibit increased mitotic activity, while the mitotic activity of outer cortical cells is suppressed (Shen *et al*., 2020b). This altered pattern of cell division contributes to the unique morphology observed in *Mtnoot* nodules that resemble that of actinorhizal nodules with central vascular tissue. *NOOT* also has a mild influence of the size of root apical meristem (RAM) in *Medicago*, with *noot* mutants displaying changes in spatial development of root cell files, although the overall impact on root architecture has not been reported (Shen et al., 2019). It is not clear if COCH/NOOT also influence the earlier stages on nodulation including rhizobial infection, as well as other key elements of root development including root architecture or interaction with arbuscular mycorrhizal symbiosis.

Plant hormones including auxin, cytokinin and gibberellin are key regulators of cell division, differentiation and organ development, including regulating lateral root and nodule development (reviewed by Azarakhsh and Lebedeva, 2023; Jan et al., 2024; Velandia *et al*., 2022). Indeed, hormones may be a key mechanism in the common and divergent developmental pathways in root and nodule development; auxin and gibberellin have positive roles in the development of both organs (Tu et al., 2024; Ubeda-Tomás et al., 2009; Velandia et al., 2024), while cytokinin promotes the development of nodules but suppresses lateral root emergence and growth (e.g. Casimiro et al., 2001; Gonzalez-Rizzo et al., 2006; Riefler et al., 2006). However, relatively little is known about how NOOT/COCH interacts with plant hormones to control nodule development. In *Medicago*, there is disruption in pattern of gibberellin (as monitored by the biosensor GIBBERELLIN PERCEPTION SENSOR 2) in root-nodule hybrids of *noot1 noot2* double mutants, in particular a reduction of gibberellin in the nodule meristem (Drapek et al., 2024). As outlined below, transcript profiling in the days following inoculation with rhizobia indicate auxin and cytokinin pathway gene expression are disrupted in *noot noot2* mutant and *NOOT/NOOT2* overexpression lines compared to wild type (Lee *et al*., 2024).

Detailed mutant and RNAseq analysis have indicated some of the factors that may interact with *NOOT/NOOT2* to control nodule identity in *Medicago*. *NOOT/NOOT2* appears to act downstream of transcription factors *LSH1/LHS2* belonging to the LIGHT-SENSITIVE SHORT HYPOCOTYL (LSH) family that are key factors mediating the divergence of lateral roots and nodules in *Medicago* (Lee *et al*., 2024). An interaction between these pathways is supported by the fact that there is strong overlap in the genes mis-regulated in *noot noot2* and *lsh1 lsh2* double mutant roots, and *lsh1 lsh2* mutant displays at least some root-nodule hybrid structures (Lee *et al*., 2024). However, the fact that ectopic expression of *NOOT/NOOT2* could not fully rescue *lsh1 lsh2* phenotype (Lee *et al*., 2024) indicates these genes may also act in somewhat parallel roles during early nodule development. A range of auxin and cytokinin biosynthesis, transport and response proteins presented altered expression in *noot noot2* mutant and *NOOT/NOOT2* overexpression lines in the days following inoculation compared to wild type (Lee *et al*., 2024). Indeed, the pattern of hormone gene expression in *noot noot2* was similar to that of *lsh1 lsh2* mutants, leading to the hypothesis that LSH acts via NOOT to regulate hormone dynamics (Lee *et al*., 2024). An examination of auxin and cytokinin levels and response during nodule organogenesis, accompanied by tissue specific RNAseq studies, would enable us to identify nodule-specific targets acting downstream of NOOT/COCH.

Autoregulation of nodulation (AON) is a root-shoot-root systemic signalling system that limits nodule number. AON signalling is enhanced by the presence of more mature nodules (Kassaw et al., 2015; Li et al., 2009) and AON suppresses subsequent infection and nodule organogenesis (reviewed by Roy and Muller, 2022; Yoro et al., 2019). Furthermore, studies in at least in some legumes have revealed elements of the AON system (CLAVATA1, KLV, miR2111) not only influence nodule number, but also independently influence the number of lateral roots and lateral root development (Day et al., 1986; Goh et al., 2018; Lagunas et al., 2019; Wopereis et al., 2000; Zhang et al., 2021), although this did not appear to be the case in pea (Wang et al., 2020).Thus, it would be interesting to examine if root-nodule hybrids of *coch/noot* mutants interact with the AON system in a similar way to wild type and if elements of AON may influence the development of root-nodule hybrids in *coch*.

In this study, the interaction between *PsCOCH* and plant hormones auxin, cytokinin and gibberellin in nodule development was examined in detail. This included examination of auxin and cytokinin response using hormone-responsive promoter::*GUS* studies, hormone measurements and tissue specific RNAseq in wild type and *coch* mutants. We found that *PsCOCH* is required for spatial tight regulation of auxin and cytokinin during nodule organogenesis and identify key hormone and signalling genes regulated by *COCH* during nodule development. We also found that *PsCOCH* in the root appears to act specifically in nodule organogenesis, with no obvious effect on rhizobial infection, root architecture or consistent effect on colonisation by arbuscular mycorrhizal fungi.

## Results

### *PsCOCH* impacts on nodulation, root development and arbuscular mycorrhizal symbioses

As the impact of *coch*/*noot*/*nbc1* mutations on infection thread development or arbuscular mycorrhizal symbioses has not been reported in any species this was characterised in *coch-JI2165, coch-JI2757* and corresponding wild type lines. Mutation of the *PsCOCH* gene did not result in any change in number of infection threads (Figure 1A) and no consistent effect of *coch-JI2165* mutation on infection thread number was observed across multiple independent experiments (data not shown). *coch-JI2165* mutant lines developed a range of mature nodulation phenotypes, including approximately 50% relatively normal nodules and approximately 50% abnormal nodule types not seen in wild type lines (Figure 1B), including nodules bearing root-like structures at tips (Figure 1C) and heart-shaped nodules with central vascular system (with a clear invagination at nodule centre; Figure 1D to 1E), consistent with other pea *coch* lines (Ferguson and Reid, 2005; Liu *et al*., 2023). Sections taken through nodules revealed that abnormal *coch-JI2165* nodules have reduced number of vascular bundles (3 instead of 4 presents in single lobed WT nodules) and these strands are thickened (Figure 1D and 1F). The abnormal *coch* nodules have additional uninfected cell layers in the outer cortex and the intermediate and nitrogen fixation zones of *coch* abnormal nodules contain less rhizobia than wild type nodules (Figure 1D and 1E), indicating defects in bacterial colonization and symbiosome formation.

**Figure 1.**
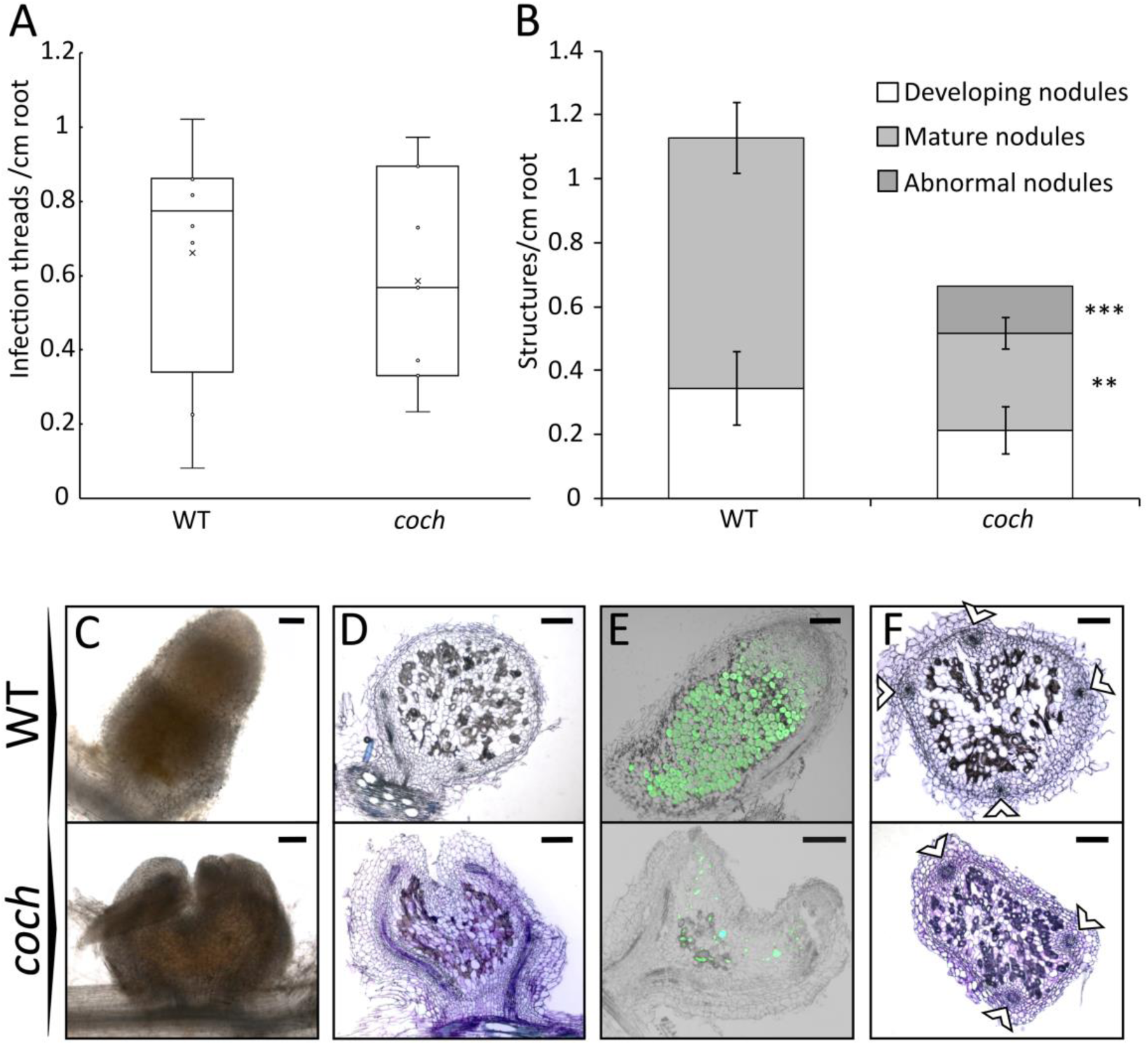
Nodule development defects in the *coch* mutant include root-like structures, reduced colonization, and vascular disorganization. Nodulation phenotypes of wild-type Weitor (WT) and *coch*-JI2165 mutant plants at 6 weeks old, 4 weeks after rhizobial inoculation (A) Number of infection threads per cm of root length. Boxes represent the interquartile range, horizontal lines show the median, and whiskers indicate the full data range. (B) Number of nodules per cm of root length. Data in (A–B) represent mean ± standard error (SE), *n* = 7. Asterisks indicate statistical significance based on *t-test*: \*\**p* < 0.01, \*\*\**p* < 0.001. (C) Brightfield microscopy images of typical nodules in wild-type and *coch* mutant plants, showing an abnormal nodule with a root-like structure at the tip in the mutant. (D) Longitudinal sections of wild-type and *coch* mutant nodules, showing the characteristic heart-shaped morphology in the mutant. (E) Brightfield and GFP-fluorescence merged images of longitudinal sections showing GFP-tagged *Rhizobium leguminosarum* within nodules. (F) Transverse sections of wild-type and *coch* mutant nodules under brightfield microscopy, with white arrows indicating vascular bundles. Scale bars = 200 µm. (D, F) Sections were stained with toluidine blue.

Overall, the *coch-JI2165* mutation did not have a consistent effect on the number of developing or mature nodules (Figure 1B; data not shown). There was also no consistent effect of *coch* mutations on root architecture in plants grown under sterile conditions (Table S1), consistent with previous reports of other *coch* alleles (Ferguson and Reid, 2005). We examined the influence of two *coch* mutant alleles on arbuscular mycorrhizal symbiosis. A small but significant increase in the amount of the root colonised by arbuscular mycorrhizal fungi, including all fungal structures and arbuscules, was observed in *coch-JI2165* plants compared to respective wild type plants, although no change was observed in colonisation rates in *coch-JI2757* lines (Figure S1).

The regulation of nodule number is under strict control of the systemic autoregulation of nodulation (AON) pathway (reviewed by Roy and Muller, 2022) and in pea this negative feedback loop is activated by nodule primordia development (Li *et al*., 2009). To explore if the formation of abnormal *coch* nodules is regulated by the AON pathway, the phenotypes of double mutants disrupted in COCH and key elements of the AON pathway, CLE peptide receptors *Ps*NARK (CLV1 orthologue) and *Ps*CLV2, were examined (Figure 2). As previously reported, *Psnark* and *Psclv2* single mutants display significant increase in nodule number (e.g. Sagan and Duc, 1996), and a large increase in total nodule number was also observed in *Psnark coch* and *Psclv2 coch* double mutants compared to wild type lines (Figure 2A). The *coch* mutation did not have a consistent influence on total nodule number in different AON mutant backgrounds. However, the proportion of abnormal nodules was actually enhanced in both *Psnark coch* and *Psclv2 coch* compared to *coch* single mutant plants, rising from approximately 70% abnormal nodules in *coch* mutants to approximately 85-95% abnormal nodules in AON mutant backgrounds. Several genes involved in the AON pathway showed differential expression in *coch* mutant nodules (Figure S2D). The expression of the putative CLE receptor genes *KLV1* and *KLV2* were upregulated in *coch* developing nodules, whereas their expression remained unchanged in wild-type nodules compared to root primordia expression. In contrast, the expression of *TML2*, a transcription factor that may act downstream of AON pathway to suppresses nodulation (Gautrat et al., 2019; Lebedeva et al., 2022; Nishida et al., 2016), increased in wild-type mature nodules but remained unchanged in *coch* mature nodule tissue, suggesting that AON pathway activation may be at least partially dependent on *COCH* function.

**Figure 2.**
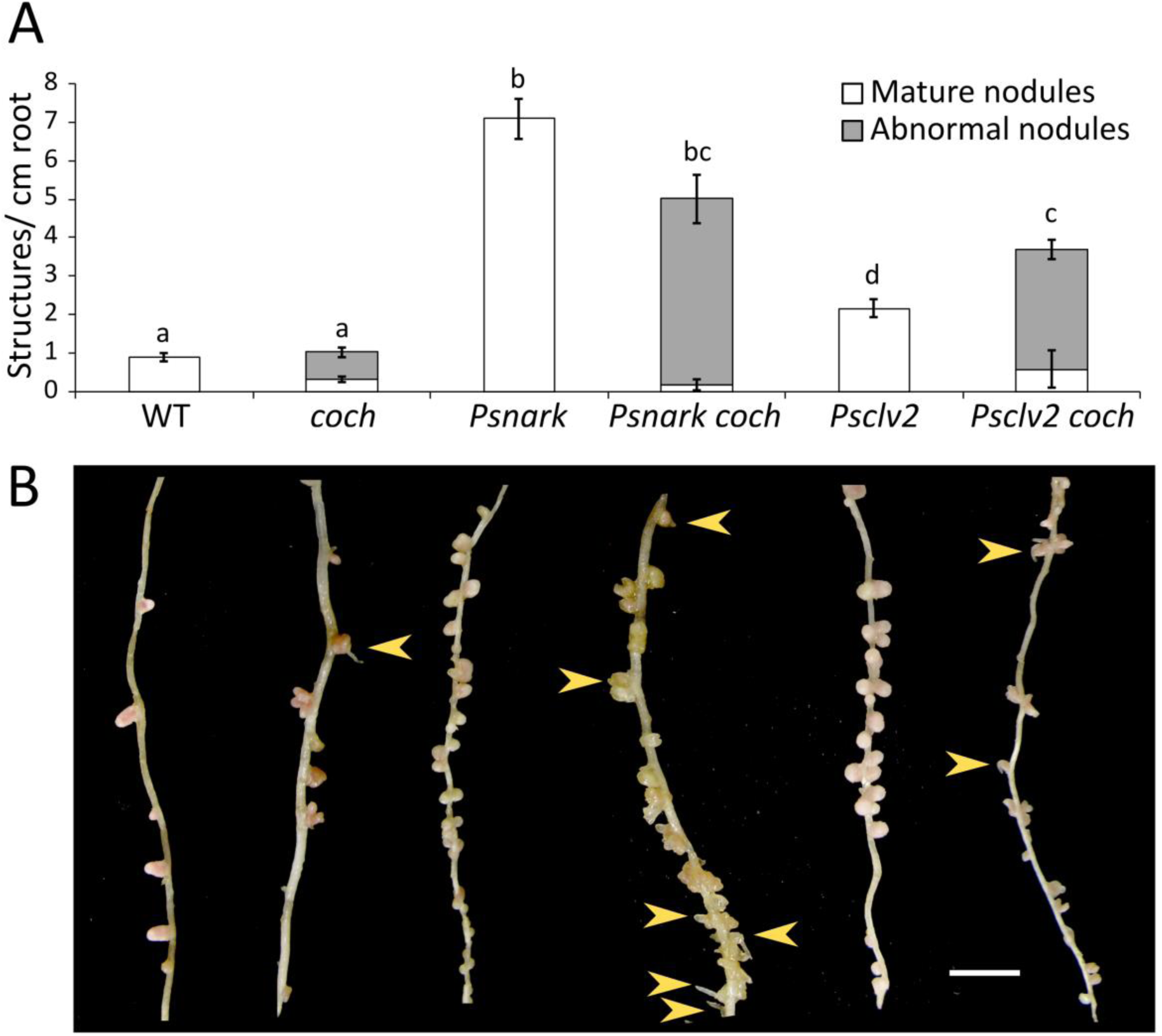
Double *coch*-AON mutants develop a higher proportion of abnormal nodules compared to single mutants and wild type. (A) Number of mature, normal, and abnormal nodules per cm of root in wild-type plants, single mutants, and F3 double mutant segregants from a cross between *coch-JI2165* and *Psnark Psclv2*. Data are shown as mean ± SE, *n* = 5–6. Different letters indicate statistically significant differences in total nodule number (mature + abnormal), assessed using Tukey’s Honestly Significant Difference (HSD) test (*p* < 0.05). (B) Representative root images showing typical secondary roots. Yellow arrows indicate abnormal nodules displaying root-like outgrowths at the tip. Scale bar = 1 cm. Plants were grown for 4 weeks, and nodulation phenotypes were assessed 3 weeks post-inoculation with *Rhizobium*.

### RNAseq analysis of roots and nodules in wild type and *Pscoch* mutants

Previous RNA-seq in *Medicago* investigated early expression profiles in *noot1/noot2* (within 72 h post inoculation; Lee *et al*., 2024) but given *coch* largely acts during nodule organogenesis, we performed tissue specific RNA-seq analysis on root primordia, developing nodules, and mature nodules of wild-type and *coch*. Root primordia expression profiles were largely similar between genotypes, with only 23 differentially expressed genes (DEGs) identified, and the samples clustered closely in the principal component analysis (PCA) (Figure 3A). However, developing *coch* nodules a more root-like transcript profile, clustering more closely with wild-type root primordia than with wild-type developing nodules (Figure 3A). Indeed, *coch* developing nodules showed roughly half as many differentially expressed genes as wild-type developing nodules. (Figure 3A), indicating a divergence in expression profile in early nodule development between *coch* and wild type. The transcript profile of mature nodules of wild-type and *coch* overlapped and were distinct from developing nodules, suggesting the gene expression patterns in mature nodules converge in the two genotypes.

**Figure 3.**
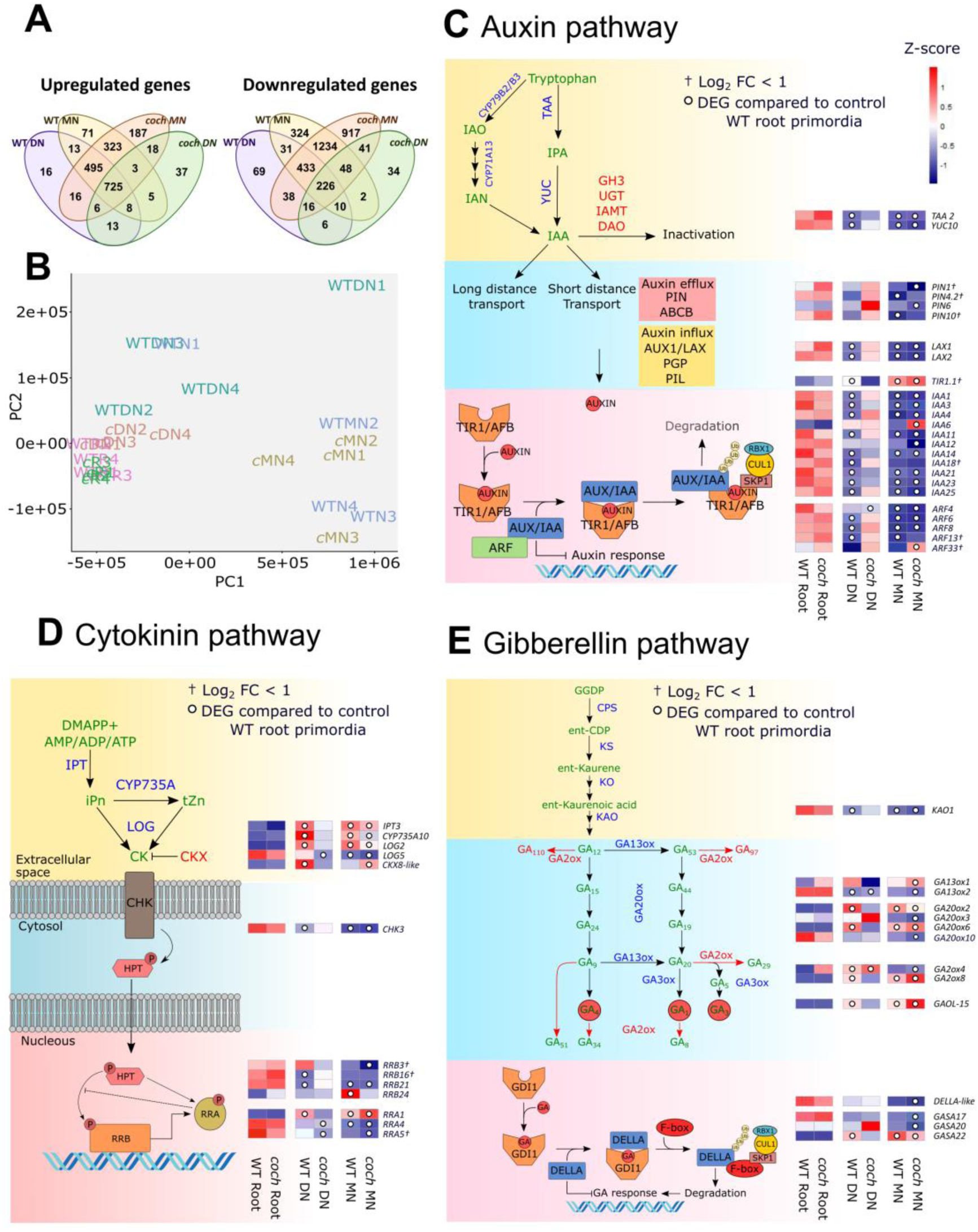
Transcriptomic comparison of wild-type and *coch* mutant nodules reveals hormone pathway alterations. (A) Venn diagram showing the overlap of differentially expressed genes (DEGs) that were upregulated or downregulated in wild-type and *coch* mutants in developing and mature nodules. (B) Principal component analysis (PCA) of gene expression profiles across samples. Each point represents a biological replicate (n = 4). (C–E) Simplified hormone pathway diagrams and heatmaps showing expression patterns of genes involved in auxin (C), cytokinin (D), and gibberellin (E) biosynthesis, signalling, and catabolism. Hormone species and intermediates are shown in green, biosynthesis enzymes in blue, and catabolic enzymes in red. Arrows indicate enzymatic reaction directionality. Only genes showing a differential expression pattern between wild-type and *coch* mutants are included in the heatmaps. The heatmap colour scale represents Z-scores of normalized gene expression. White circles within each heatmap cell indicate that the gene in that specific condition was significantly differentially expressed relative to the wild-type root primordia (q-value < 0.05). Genes marked with a cross (†) indicate that although differentially expressed, their absolute log₂ fold change (log₂FC) was < 1. Samples were collected from 6-week-old plants, 4 weeks post-inoculation with *Rhizobium*. Tissues included root tips, developing nodules, and mature nodules for wild-type, and abnormal nodules for *coch* mutants. Abbreviations: DMAAP⁺: N6-(Δ2-isopentenyl)adenosine monophosphate. AMP/ADP/ATP: Adenosine mono/di/tri-phosphate. iPn: Isopentenyladenine. tZn: *Trans*-zeatin. CK: Cytokinin. IAO: Indole-3-acetaldoxime. IAN: Indole-3-acetonitrile. IPA: Indole-3-pyruvic acid. IAA: Indole-3-acetic acid. GGDP: Geranylgeranyl diphosphate. *ent-CDP*: *ent*-Copalyl diphosphate. GA: Gibberellin. DN: Developing nodule. MN: Mature nodule. Gene symbols and full names are listed in Table S3.

To gain insight into the biological processes affected by the *coch* mutation, we performed Gene Ontology (GO) enrichment analysis on the subset of genes that were differentially expressed in wild-type developing and/or mature nodules but not differentially expressed in *coch* mutant developing or mature nodules, relative to wild-type root primordia (Figure S2A-B). Genes uniquely upregulated in wild-type nodules were enriched with GO terms predominantly associated with energy and carbon metabolism, including key components of the tricarboxylic acid (TCA) cycle and carbohydrate-derived energy production (Figure S2A). Other enriched GO categories included lipid metabolism pathways, such as sphingolipid, isoprenoid, and ubiquinone biosynthesis; pathways important for formation of the symbiosome, (Moore et al., 2021) and mitochondrial oxidative energy metabolism to sustain nitrogen fixation (Sin et al., 2024) respectively. Conversely, genes uniquely downregulated in wild-type nodules were enriched for GO terms related to plant defence, immune responses, and reactive oxygen species signalling (Figure S2B). This group included multiple leucine-rich repeat (LRR) resistance genes involved in effector triggered immunity (ETI) (Tong et al., 2023), and peroxidases. ETI responses are thought to be repressed during nodule development with some LRR proteins contributing to host specificity (Gourion et al., 2015; Yang et al., 2010). This suggests a role for COCH in suppressing defence responses to facilitate symbiosis. Notably, two genes putatively involved in root cap formation and stem cell differentiation in the root meristem also appeared to be downregulated in a *COCH*-dependent manner.

To investigate the genes potentially mis-regulated in *coch* that may contribute to the formation of chimeric nodule–root structures, we investigated the expression of genes previously characterized as regulators of root organogenesis (Figure S2C). Several genes that were differentially expressed in wild-type developing and mature nodules compared to wild type root primordia showed no significant changes in the corresponding *coch* mutant tissues. Notably, components of the auxin signalling pathway, including *ARF6/8* and the auxin influx transporter *AUX1*, were downregulated during nodule development in the wild type but remained unchanged in *coch* developing nodules and more closely resembled that of root primordia (Figure S2C). In addition, several key root development transcription factors (TFs) were differentially expressed in wild-type nodules but not in the *coch* mutant. Specifically, *coch* mutants showed reduced expression of *PLETHORA3* (*PLT3*) in developing nodules and *PLT5* in mature nodules compared to the corresponding wild-type tissues. PLT are known to orchestrate lateral root growth and differentiation of the emerging root layers (Du and Scheres, 2017; Santuari et al., 2016). Other important TFs involved in root and nodule development failed to be down-regulated in *coch* nodules (Figure S2C), including *HB2*, *GRF6* and members of the *KNOX* family (Dean et al., 2004; He et al., 2020; Qi and Zheng, 2013; Rodriguez et al., 2015; Yan et al., 2024). Conversely, LOB DOMAIN CONTAINING PROTEIN *LBD11*expression was reduced in *coch* mature nodules. LBD11, a homologue of *Arabidopsis thaliana* LBD3, functions in secondary root growth downstream of cytokinin signalling (Ye et al., 2021) and its accumulation has been shown to inhibit lateral root formation (Zuo et al., 2023). Other genes showing differential expression between wild-type and *coch* nodules were associated with cell cycle regulation, cell wall modification, and epigenetic control (Figure S2C).

Genes related to early symbiotic signalling, rhizobial infection, nodule organogenesis, and nodule function displayed a delayed transcriptional response in *coch* developing nodules (Figure S2D). These genes were mis-regulated in developing *coch* nodules but appeared to recover to expression levels similar to wild type in mature nodules, suggesting a temporal delay in the activation of the nodulation program in the *coch* plants. The expression of *COCH1* and *COCH2* was not significantly different between wild-type and *coch* mutant nodules, indicating that *COCH2* is not under feedback control by *COCH1*. Expression patterns of genes related to auxin, cytokinin, and gibberellin signalling are discussed below.

### Constitutive overexpression of *PsCOCH* can induce root-nodule hybrids

The effect of the loss of function of COCH/NOOT1/NBC1 is well understood across several species but the impact of ectopic expression of COCH on nodulation is unknown. The full length *PsCOCH* gene was expressed under both tissue-specific and constitutive promoter in wilt type pea hairy roots (Figure 4). In pea roots, the *35S* promoter is constitutively expressed in all root tissue, *AtCO2* promoter is expressed in basal meristematic zone as seen in *Aarbidopsis* (Figure S3; Heidstra et al., 2004) and the *AtCASP* promoter is expressed in endodermis (Velandia *et al*., 2024). *CASP::GUS* was used as a control. Quantitative real-time PCR analysis of whole root samples confirmed that *PsCOCH* expression was significantly elevated in *35S::COCH* roots compared to *CASP::GUS* controls, while modest increases were also observed in *CASP::COCH* and *CO2::COCH* roots (Figure S4).

**Figure 4.**
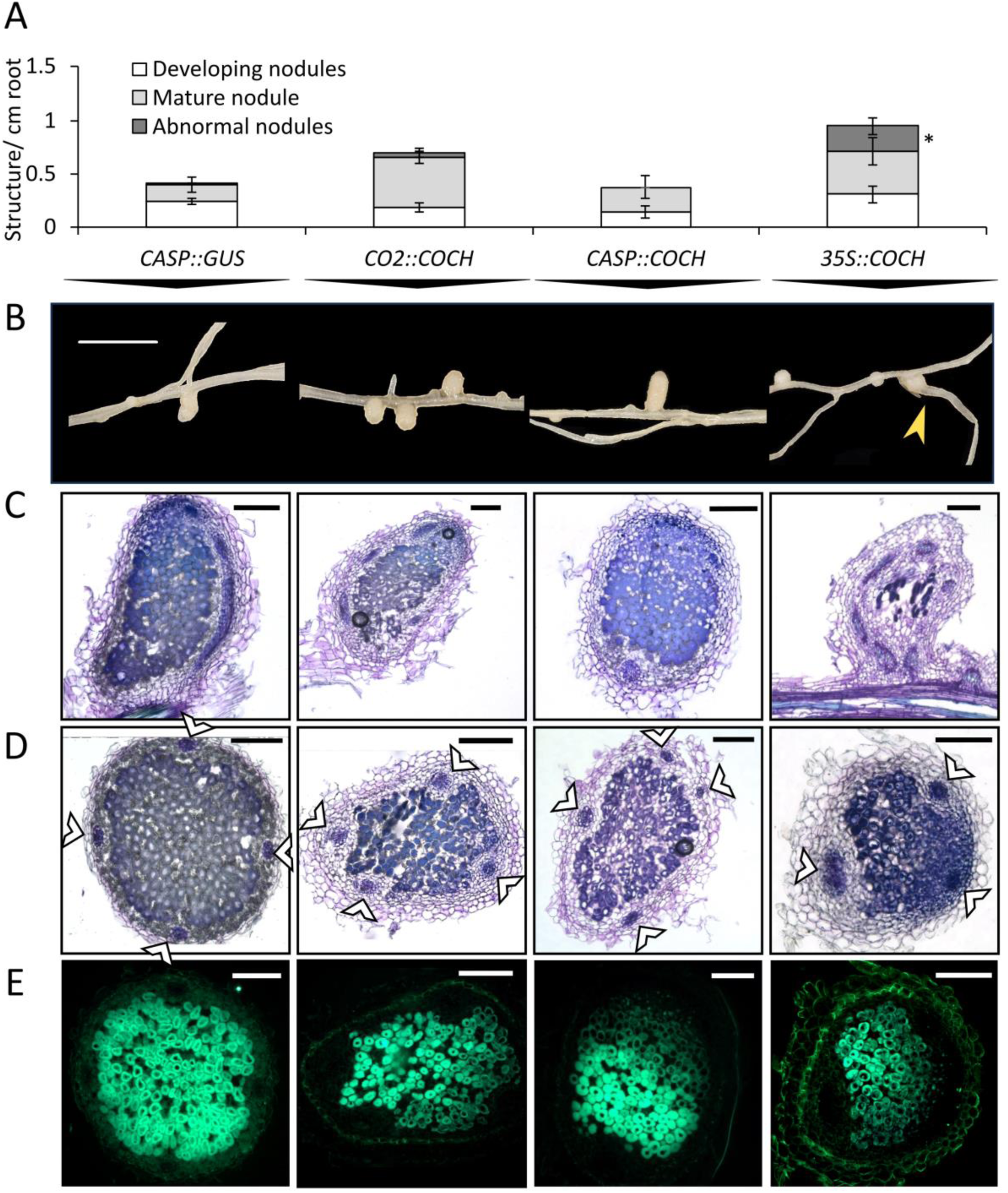
Ectopic expression of *PsCOCH* in hairy roots induces abnormal nodules and nodule–root hybrids in wild-type *Parvus*. Nodulation phenotypes were assessed 3 weeks post-inoculation in hairy root-transformed wild-type *Parvus* plants expressing *PsCOCH* under a constitutive promoter (*35S::COCH*), an endodermis-specific promoter (*CASP::COCH*), or a meristem-specific promoter (*CO2::COCH*). Plants transformed with *CASP::GUS* (endodermis-specific promoter driving β-glucuronidase) were used as controls. (A) Number of developing, mature, and abnormal nodules per cm of root. Data represent mean ± SE, *n* = 8–9. *p* < 0.05 (t-test). (B) Images of root segments showing representative nodules from each transformation condition. Scale bar = 2 cm. (C) Longitudinal sections of representative nodules from each treatment stained with toluidine blue. (D–E) Transverse sections of representative nodules stained with toluidine blue, shown in brightfield (D) and GFP fluorescence (E). GFP indicates *Rhizobium leguminosarum*. White arrows (D) indicate vascular bundles. Scale bars = 200 µm.

Expression of *PsCOCH* under *CO2* promoter or *CASP* promoter had no observable effect on nodule number or development compared to the *CASP::GUS* control (Figure 4A). Strikingly, *35S::COCH* overexpression resulted the production of abnormal nodules exhibiting characteristics similar to those of *coch* mutants. These included nodules with irregular shapes (Figure 4C), thickened vascular strands, a reduced number of vascular strands (Figure 4D), and root-nodule hybrids (Figure 4B). No changes in root architecture or vascular patterning were observed in any of the constructs (data not shown).

### Elevated cytokinin levels and ectopic cytokinin response in *Pscoch* nodules

Cytokinin is essential for nodule organogenesis; nodules develop at the site of cytokinin foci that form due to tight spatial regulation of cytokinin biosynthesis and catabolism (reviwed by Nadzieja et al., 2019; Velandia *et al*., 2022). This positive role for cytokinin in nodule organogenesis is contrasted with a negative role in lateral root development (Azarakhsh and Lebedeva, 2023; Laplaze et al., 2007; Reid et al., 2017). It has been proposed that this specific role for cytokinin in nodule organogenesis may be due to the key nodulation gene *NIN* upregulating cytokinin production that in turn promotes auxin biosynthesis via LBD16, STY and YUC to activate nodule organogenesis (Schiessl *et al*., 2019).

We monitored endogenous cytokinin levels in mature nodules of wild type and *coch* mutant. The level of cytokinin trans-zeatin riboside (t-ZR) was significantly higher in *coch* mature nodules than levels found in mature wild type nodules (Figure 5A). Cytokinin response was also assessed using the *TCSn::GUS* reporter. In wild-type roots, *TCSn::GUS* activity was typically observed in cells surrounding infection threads connected to nodules and in most developing nodule primordia, as observed previously in pea (Figure 5B) (Velandia *et al*., 2024). In wild type mature nodules *TCSn::GUS* signal is largely confined to the nodule meristem and has a faint presence in the vasculature (Figure 5F, 5I). Quantification of GUS presence indicated that 60–70% of wild-type infection threads, developing primordia, and mature nodules exhibited a detectable *TCSn::GUS* signal (Figure 5B). A different pattern of cytokinin response was observed in *coch* mutants. A significantly lower proportion of infection threads in *coch* mutants (approximately 20%) were associated with *TCSn::GUS* activity, while 100% of *coch* developing nodule primordia showed *TCSn::GUS* signal, significantly higher than observed in wild type lines (Figure 5B). Approximately 80-90% of *coch* nodules displayed *TCSn::GUS* signal, which was not significantly different to wild type (Figure 5B). However, ectopic or expanded cytokinin signalling were apparent in abnormal *coch* nodules including *TCSn::GUS* signal in thickened peripheral and central vascular strands and strong *TCSn::GUS* in the tips of heart shaped nodules and root-like structures subtending these abnormal nodules (Figure 5E-K). *TCSn::GUS* was expressed in the root tips of both wild type and *coch* mutants in a similar pattern (Figure 5L-M).

**Fig. 5.**
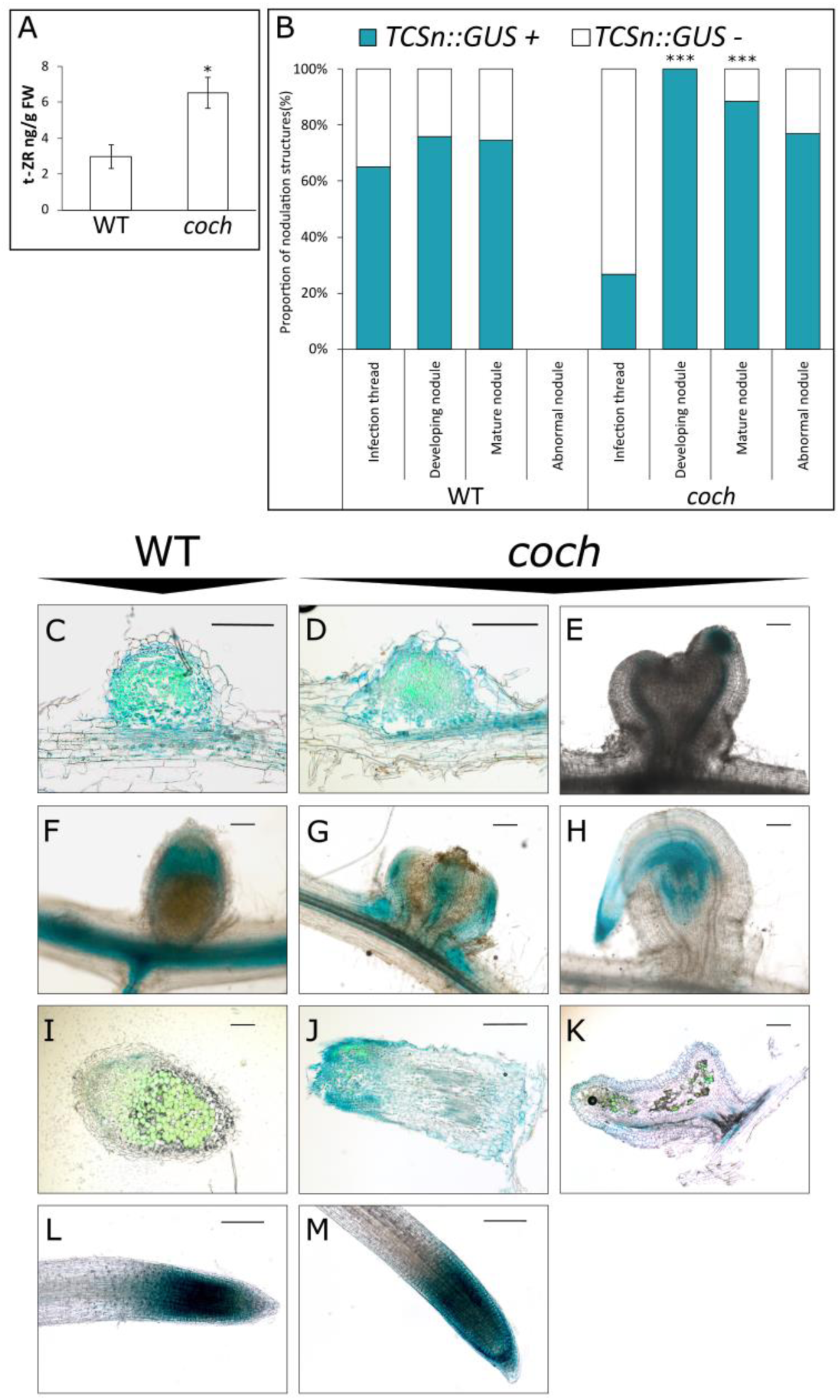
Cytokinin response and levels are elevated in *coch* abnormal nodules. Cytokinin accumulation and cytokinin-responsive promoter activity were assessed in *coch-JI2165* and wild-type *Weitor* (WT) roots transformed via hairy root transformation with the cytokinin reporter construct *TCSn::GUS* (a synthetic cytokinin-responsive promoter driving *β-*glucuronidase expression). (A) Quantification of *trans*-zeatin riboside (t-ZR) levels in mature nodules of WT and *coch*. Data are mean ± SE, *n* = 4. Asterisk indicates significant difference (*p* < 0.05, *t*-test). (B) Proportion of infection and nodule structures showing *TCSn::GUS* activity (*TCSn::GUS+*) or no detectable activity (*TCSn::GUS−*) in WT and *coch*. Chi-squared test was performed for each structure type; \*\*\**p* < 0.001, *n* = 6–7. (C–M) Representative microscopy images of *TCSn::GUS* expression in various tissues. Blue indicates GUS activity; GFP fluorescence marks *Rhizobium leguminosarum* colonization. (C–D) Longitudinal sections of developing nodules (E–H) Whole mature nodules (I–J) Longitudinal sections of mature WT nodule and abnormal *coch* nodule (L–M) Root tips expressing. Scale bars = 200 µm for all panels. Roots were sampled from 4 weeks post-*Rhizobium* inoculation.

We analysed the expression of genes involved in cytokinin biosynthesis, catabolism, and signalling in the RNA-seq dataset. The developing nodules of *coch* plants exhibited an expression profile for cytokinin biosynthesis genes *IPT3*, *LOG2*, and *CYP735A10* that is more similar to root primordia, unlike the high expression in these genes observed in developing wild type nodules (Figure 3C). In addition, several type-A and type-B cytokinin response regulators also showed altered expression in *coch* developing and mature nodules (Figure 3C), indicating that cytokinin signalling is altered in the *coch* mutant background.

### Auxin levels in *Pscoch* mutant nodules are low and the pattern of auxin response is severely altered compared to wild type

The formation of auxin maxima at the site of nodule organogenesis is achieved through precise regulation of auxin transport and/or auxin biosynthesis (reviewed by Velandia *et al*., 2022). Indeed, key cross over points in lateral root and nodule development programs involves the activation of LBD16 that in turn promotes auxin biosynthesis genes *STY* and/or *YUC* (Schiessl *et al*., 2019; Shrestha et al., 2021; Soyano *et al*., 2019). Auxin accumulation appears to occur downstream of CK dependent activation of NIN and flavonoid synthesis (Ng et al., 2015; Schiessl *et al*., 2019; Suzaki et al., 2012). Given that cytokinin levels are elevated and cytokinin response is mis-regulated in nodules of *coch* mutant plants (Figure 5), we investigated if this might lead to abnormal auxin gene expression, auxin biosynthesis and/or auxin response patterning in *coch* nodules (Figure 6).

**Fig. 6.**
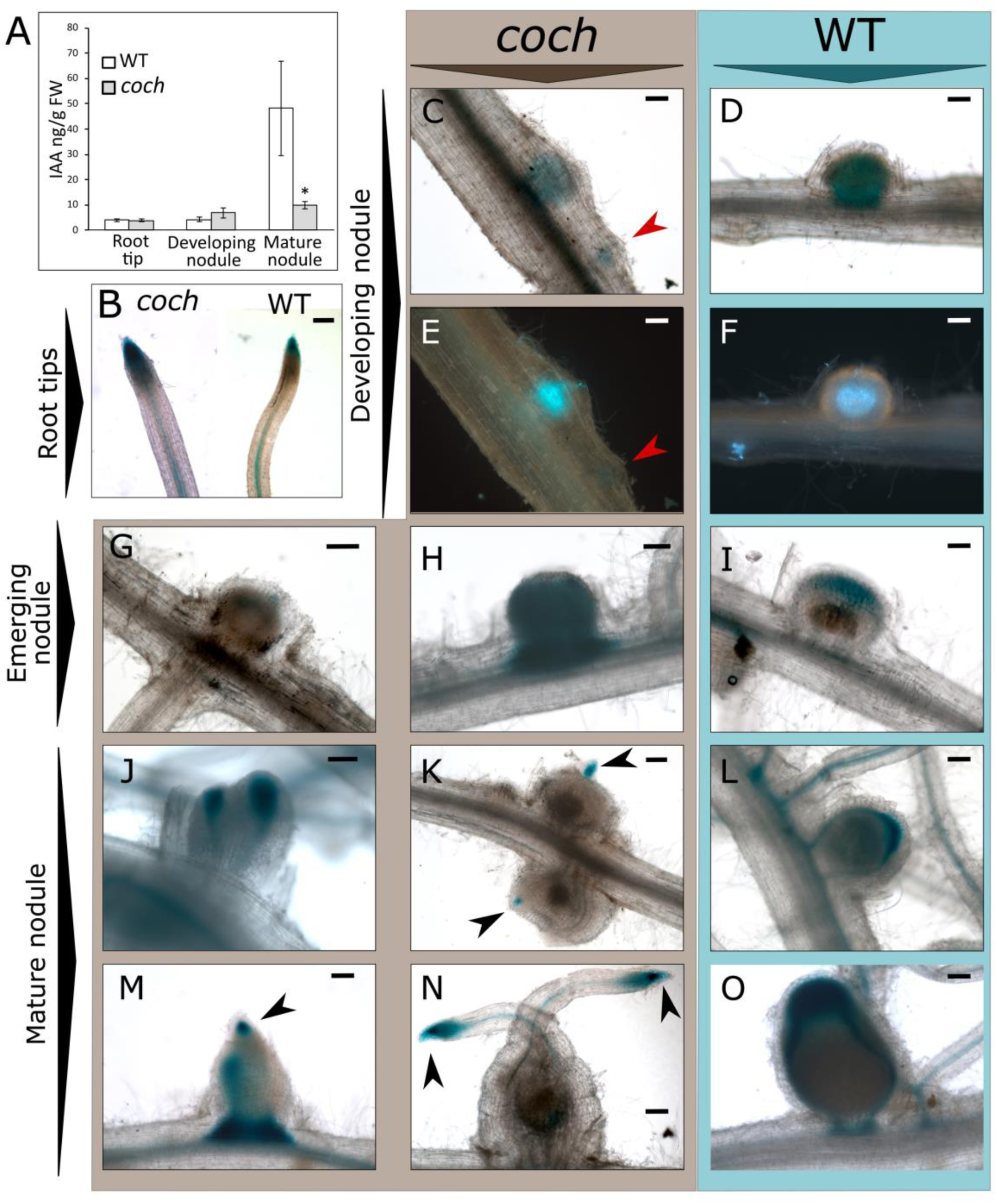
Auxin levels are reduced, and auxin response is altered in *coch* abnormal nodules. Auxin accumulation and auxin-responsive promoter activity were evaluated in *coch-JI2165* mutants and wild-type *Weitor* (WT) roots transformed via hairy root transformation with the DR5::GUS reporter construct (a synthetic auxin-responsive promoter driving *β*-glucuronidase expression). (A) Indole-3-acetic acid (IAA) content (ng/g fresh weight) in root tips, developing nodules, and mature nodules of WT and *coch*. Data are shown as mean ± SE, *n* = 4; *p* < 0.05 (Student’s *t*-test). (B–O) Microscopy images showing DR5::GUS activity in root tissues of WT and *coch-JI2165*. Blue staining indicates GUS activity (auxin response), green fluorescence indicates *Rhizobium leguminosarum* expressing GFP. (B) Representative root tips (C–F) Developing nodules in brightfield (C–D) and GFP fluorescence (E–F); red arrows indicate root primordia (G–H) Older developing nodules (J–O) Mature nodules; black arrows indicate root-like structures at the tips of abnormal *coch* nodules Scale bars = 200 µm for all panels. Tissues were sampled 4 weeks after *Rhizobium* inoculation.

Overall, the auxin response gene expression profile in *coch* developing nodules was up-regulated compared to wild type developing nodules and more closely resembled that found in root primordia (Figure 3C). In contrast to wild type nodules, in *coch* developing nodules a large number of *IAA* genes—encoding auxin response repressors—failed to be downregulated. Similarly, the expression of four *ARF* (Auxin Response Factor) genes in *coch* developing nodules was the same as seen in root primordia, in contrast to the low expression in wild type (Figure 3B). Auxin content (measured via LC-MS) and auxin response patterns (monitored using the *DR5::GUS* reporter) were assessed in nodules and lateral roots of both *coch* mutant and wild type plants (Figure 6). Auxin levels in root tips and developing nodules were comparable between *coch* and wild type, but mature *coch* nodules contained significantly lower auxin levels than mature wild type nodules (Figure 6A). Wild type and *coch* mutants expressed *DR5::GUS* throughout the developing nodule primordia and also in young root vasculature, lateral root primordia and root tips (Figure 6 B-F, Velandia *et al*., 2024). As wild type nodules developed, *DR5::GUS* expression became restricted to the nodule vasculature and nodule meristem (Figure 6 I, L, O). However, as the *coch* nodules matured two distinct patterns of *DR5::GUS* were observed. In *coch* nodules with root-like growths from tip, *DR5::GUS* signal was only found in root-like tissues, including vascular tissue and meristem, with no *DR5::GUS* signal in peripheral nodule vasculature or nodule meristem (Figure 6 K, N). Other patterns of *DR5::GUS* staining were also observed in some maturing *coch* nodules; very strong auxin response throughout the whole nodule that included cells containing bacteria (Figure 5H, Figure S5) and in heart shaped mature *coch* nodules strong auxin response in the abnormal central vascular tissue (Figure 5J, M).

### *PsCOCH* does not act via gibberellin to supress root identity in nodules

In addition to key roles for auxin and cytokinin, gibberellin is a key promoter of nodule and root development, while suppressing rhizobial infection (Ramon et al., 2021; Velandia *et al*., 2024). Studies in pea suggest gibberellin acts upstream of cytokinin to suppress infection thread progression and branching and also upstream of auxin in nodule primordia development (Velandia *et al*., 2024). Given the altered auxin and cytokinin response in of *coch* mutants (Figures 2, 5 and 6), we examined the infection and nodulation phenotypes of *coch na* double mutant plants compared to respective single mutant parents and wild-type line (Figure 7). These plants carry disruptions in both *COCH* and *NA*, a gibberellin biosynthesis gene and these lines have very low levels of active gibberellin.

**Fig 7.**
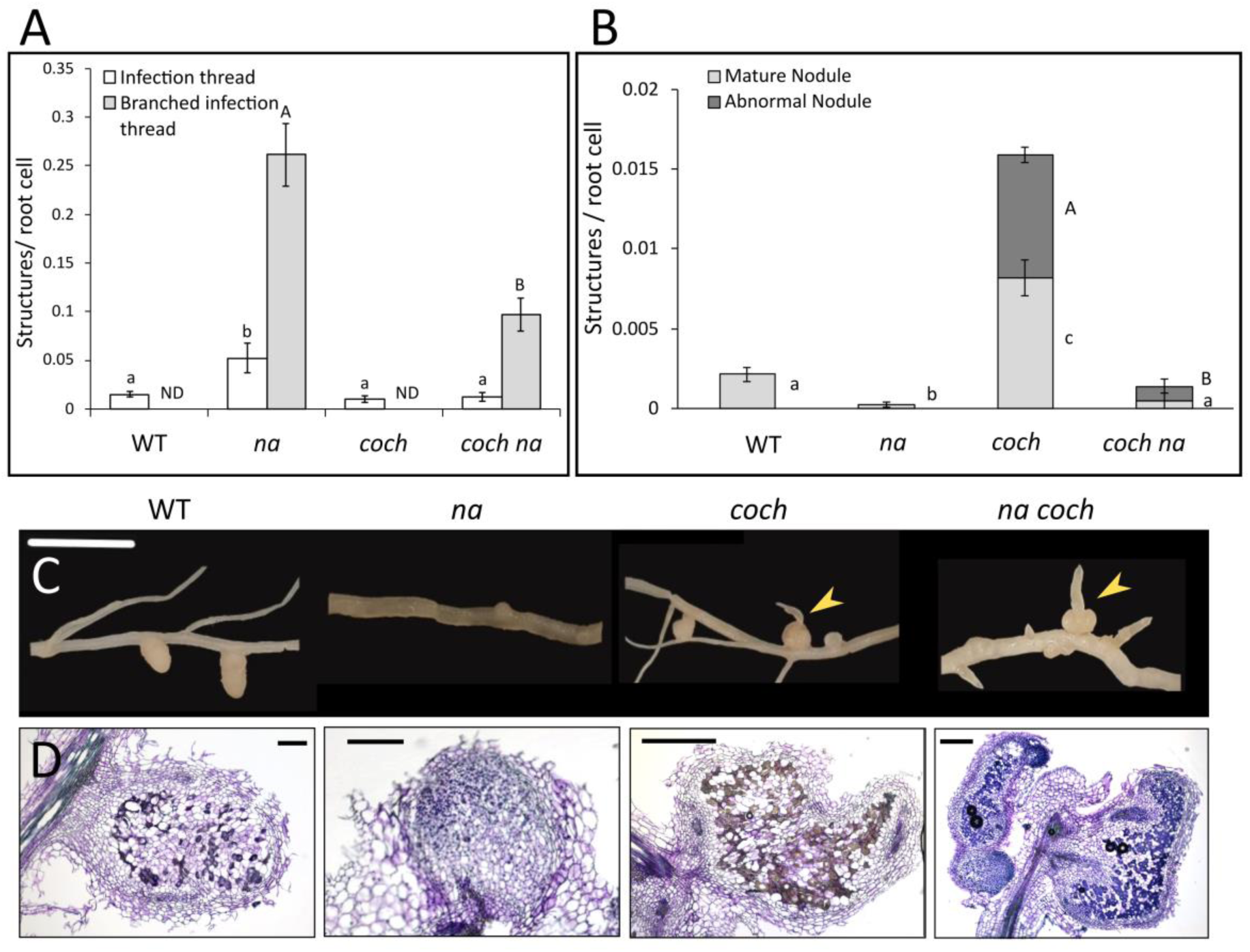
Gibberellin is not required for the formation of root-nodule abnormal structures in *coch* mutants. (A-B) Nodulation phenotypes of four weeks old F_3_ segregants from cross between *coch-JI2165* and gibberellin deficient *na* mutant, evaluated 3 weeks post inoculation. (A) Number of infection threads and branched infection threads (B) Number of normal and abnormal nodules per root cell, bars are mean ± SE, *n* = 5-6. Within a parameter, values with different letters are significantly different *p* < 0.05. ND not detected. (C) Photo of typical secondary root, yellow arrows indicate abnormal nodules with clear roots at tip. Scale bar is 2cm. (D) Longitudinal sections of nodules stained with tolidine blue; scale bar is 200µm.

As previously observed, gibberellin-deficient *na* mutants develop excessive number of infection threads, including highly branched infection threads and develop very few nodules that are underdeveloped (Figure 7A; McAdam et al., 2018; Velandia *et al*., 2024). Interestingly, double mutant *coch na* plants develop significantly less infection threads and branched infection threads than *na* (Figure 7A). As outlined above, the excessive infection thread development and branching in *na* mutants was correlated with overactivation of cytokinin response (Velandia *et al*., 2024). Gibberellin is important for nodule maturation, as both *na* and *coch na* mutant plants develop very few mature nodules compared to wild type plants (Figure 7B). However, like observed in *coch* single mutants approximately 50% of mature nodules that did form in *coch na* double mutants displayed characteristic abnormal phenotype, including nodules with roots (Figure 7B, C). Therefore, gibberellin is not required for the development of root-nodule hybrids and suggest gibberellin does not play a role in COCH control of nodule development. We also examined the expression of genes involved in GA biosynthesis, catabolism, and signalling in the RNA-seq dataset and found no consistent change in GA biosynthetic or catabolic gene expression in the *coch* mutant tissue (Figure 3D). However, some GA-responsive genes were expressed at lower levels in mature *coch* nodules than wild type mature nodules, including *GASA17, GASA20* and *GASA22* and the GA repressor *DELLA-like* (Figure 3D). This suggests that GA signalling may be somewhat suppressed in *coch* nodules.

## Discussion

The *COCH* gene functions as a key regulator in maintaining the developmental boundary between root and nodule identity, thereby ensuring proper nodule morphogenesis. In this study, we show that *COCH* regulates of key genes involved in cytokinin and auxin signalling, as well as genes associated with root development and nodulation in developing nodules. *COCH* controls auxin and cytokinin levels and spatial patterns of both cytokinin and auxin response during nodule development (Figure 8). In contrast, we did not find any compelling evidence that *COCH* requires gibberellin to regulate nodule formation (Figure 2D, 7). Together, these findings suggest *COCH* controls spatial hormone dynamics to specify and maintain nodule identity.

**Figure 8.**
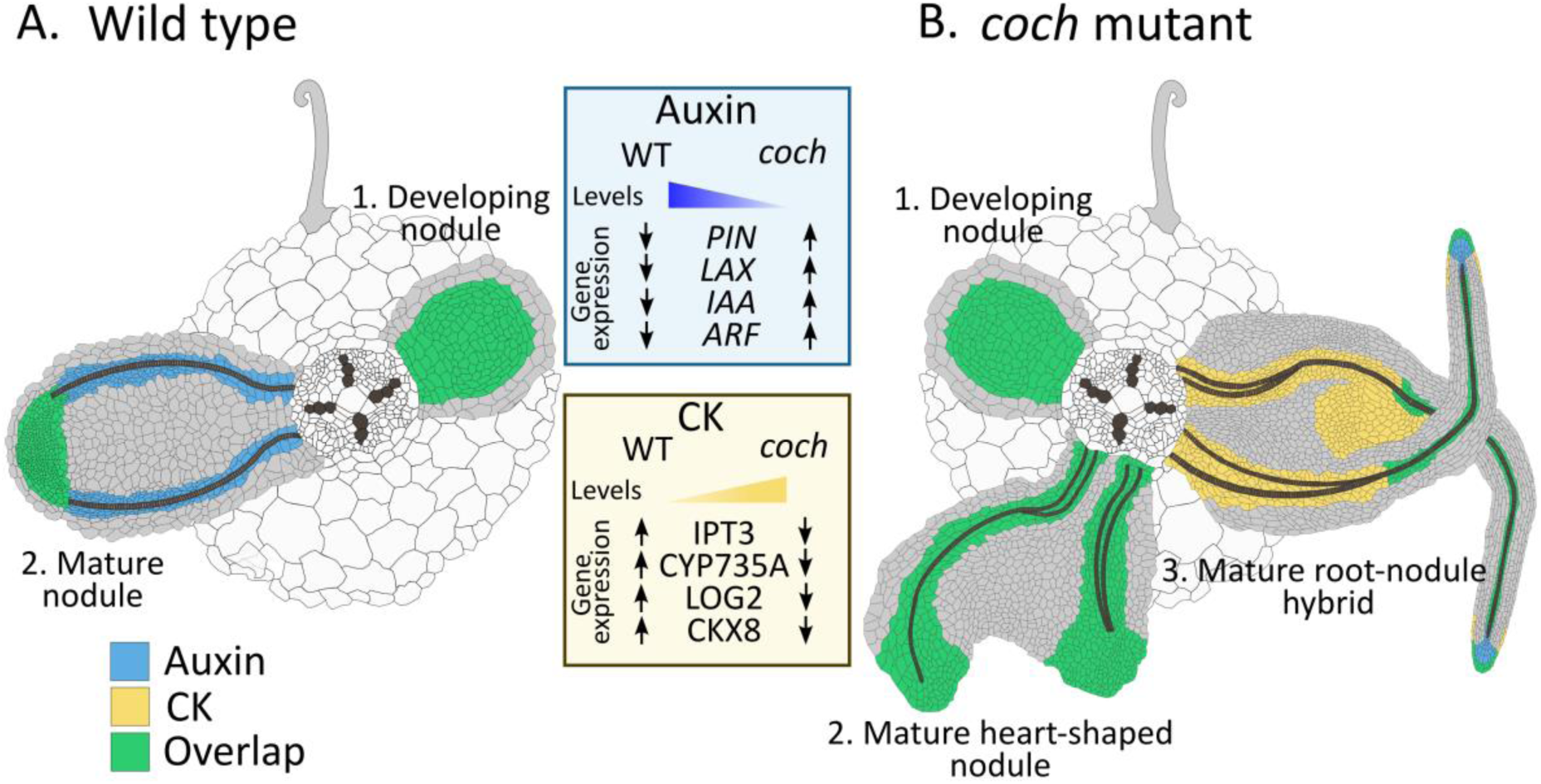
Model of hormone levels, gene expression, and spatial response of auxin and cytokinin in wild-type and *coch* mutant nodules. The spatial patterns of auxin and cytokinin response are illustrated for developing and mature nodules in wild-type and *coch* mutants. In wild-type plants, auxin and cytokinin responses overlap in the emerging nodule (A1) and remain co-localized in the meristematic region of mature nodules (A2), with auxin response also present in the nodule vasculature. In *cochleate* mutants, developing nodules initially display an overlapping auxin and cytokinin response (B1), similar to the wild type, but subsequently develop into either heart-shaped nodules (B2) or root–nodule hybrids (B3). In heart-shaped nodules, both auxin and cytokinin responses are maintained in the vasculature and at the lobe tips, but are lost from the apical meristematic region. In root–nodule hybrids, auxin response is strongly reduced and restricted to the vasculature and the tips of the fused roots, whereas cytokinin response is broadened and intensified, particularly in the abnormal vasculature and the apical zone between the merged roots. These spatial patterns are consistent with hormone quantification, which show that auxin levels are reduced in *cochleate* abnormal nodules while cytokinin levels are elevated. Gene expression analysis revealed that that auxin transport and response genes are upregulated in *coch*, and that cytokinin biosynthesis and catabolism genes are downregulated in the mutant.

*COCH* suppresses cytokinin response in the developing nodule and restricts cytokinin response to specific tissue domains during nodule maturation. The ectopic cytokinin response in *coch* mature nodules may be due to elevated cytokinin levels in this tissue compared to mature wild type nodules (Figure 5A) and/or *COCH* may target cytokinin response directly, as mature *coch* nodules displayed reduced expression of several cytokinin response regulators (Figure 2D), including *PsRRB3* that is putative ortholog of *MtRRB3* that is known to promote nodule organogenesis in *Medicago* (Tan et al., 2020). Previous studies with *coch* mutants indicate that the lateral root-like outgrowths emerge from abnormal nodule vasculature meristem which has a root-related ontogeny and is derived from pericycle cell layer divisions (Magne et al., 2018b; Shen et al., 2020a). Indeed, our studies support this as we found that the activation of cytokinin biosensor *TCSn* in abnormal developing vascular tissue and tips of root-like growths of *coch* nodules (Figure 5E, G, K) is similar to the activation of cytokinin response in pea root lateral roots (Figure 5L-M) and in the provascular cells of *Arabidopsis* emerging lateral roots (Bielach et al., 2012). *COCH* is also required for establishment of cytokinin biosynthesis and response expression patterns genes during nodule organogenesis. Both cytokinin biosynthesis gene *PsIPT3* and cytokinin receptor gene *PsCHK3*, whose orthologues in *Medicago* are known to be required for nodule development (Boivin et al., 2016; Triozzi et al., 2022), fail to be up-regulated in developing *coch* nodules. Indeed, recent studies show that the expression of *MtIPT3* is specifically induced in the pericycle during nodule primordia formation to enable cytokinin biosynthesis and activation of nodule organogenesis (Triozzi *et al*., 2022). In contrast to *Medicago*, that displayed elevated expression of cytokinin response regulator *MtRRA4* in developing nodules (Ariel et al., 2012), this was not observed for *PsRRA4* in wild type developing nodules but *PsRRA4* and *PsRRA5* were ectopically expressed in developing *coch* nodules.

Studies across many species indicate cytokinin influences auxin to control nodule organogenesis (reviewed by Velandia *et al*., 2022) and we found that regulation of cytokinin by *COCH* was correlated with auxin level and response. Dramatically different patterns of auxin response were seen between mature *coch* and wild type nodules (Figure 6, 8). Most *coch* nodules displayed auxin response only in the root-like protrusions at nodule tips and on average the mature nodules of *coch* mutants displayed much lower levels of auxin. However, there was a range of auxin response phenotypes, including a small number of maturing *coch* nodules displayed activation of auxin response throughout the nodule or in vascular strands of heart-shaped nodules. This heterogeneous reprogramming of auxin response in *coch* mutants is consistent with the range root-nodule hybrid phenotypes observed in *coch* nodules. The only major changes in auxin gene expression in mature *coch* nodules was altered expression of several auxin influx genes, including reduced expression of *PsPIN1* and *PsPIN6* and elevated expression of *PsPIN4.2* and *PsPIN10* compared to wild type mature nodules (Figure 2C). Recent work in *Medicago* has highlighted key roles for *MtPIN2*, *MtPIN4* and *MtPIN10* in establishing the nodule meristem (Xiao et al., 2025). Thus, *COCH* may regulate correct patterns of auxin transport in the maturing nodule; a key determinant of nodule initiation and development (reviewed by Velandia *et al*., 2022). Taken together, *COCH* may spatially restrict cytokinin response that in turn promotes correct auxin accumulation in vascular and nodule meristem to promote proper nodule development (Figure 8).

The *COCH* gene is essential to transcriptionally reprogram root tissue to form nodules, controlling the expression of both key hormone, root and nodulation genes. Tissue specific RNA-seq found that developing *coch* nodules displayed transcriptional profile that more closely resemble root primordia than wild type developing nodules (Figure 3B), complementing previous studies in *Medicago* that found *noot noot2* mutants failed to regulate around 40% of genes that wild type plants differentially expressed within 24-72 hours of inoculation with rhizobia (Lee *et al*., 2024). We found even larger divergences in expression between *coch* and wild type occur in the developing nodule tissue (Figure 3, Figure S2). *COCH* is required during early nodule development to suppress expression of auxin biosynthesis, transport, perception and response genes (nine *AUX/IAA* repressors, four *ARFs*, and two *AUX1/LAX* transporters). Several of these genes have homologs in other species that are known to regulate lateral root development; *ARF8* promotes lateral roots and *IAA14* suppresses lateral roots in *Arabidopsis* (Fukaki et al., 2005; Fukaki et al., 2002; Wang et al., 2015) and in soybean *GmARF8* must be locally suppressed to enable nodules to form (Wang *et al*., 2015). Notably, auxin-regulated transcription factors *PLT3* and *PLT5* involved in root apical meristem maintenance and differentiation and correct formation and patterning of lateral roots (Du and Scheres, 2017; Galinha et al., 2007; Hofhuis et al., 2013; Mähönen et al., 2014) fail to be up-regulated in *coch* developing nodules and are downregulated in mature *coch* nodules respectively. In contrast, several other genes with important roles in lateral root development *HB2*, *GRF6* and members of the *KNOX* family (Dean *et al*., 2004; He *et al*., 2020; Qi and Zheng, 2013; Rodriguez *et al*., 2015; Truernit and Haseloff, 2007; Yan *et al*., 2024), are up-regulated in *coch* developing or mature nodule compared to wild type (Figure S2C). The majority of nodulation genes upregulated in wild type developing nodules fail to be upregulated in developing *coch* nodules, although their expression is restored in mature *coch* nodules (Figure S2D). Therefore, *COCH* may act upstream of cytokinin and auxin early in nodule development to maintain nodule identify by suppressing key root identity genes and promoting key nodulation genes and this sets the organs on a nodule trajectory.

*PsCOCH* appears to act specifically in the root to influence nodule organogenesis, with no consistent effects on rhizobial infection, root architecture or colonisation with arbuscular mycorrhizal fungi (Figure 1, Figure S1, Table S2). The fact that no change in root architecture was detected in *coch* mutants is consistent with no major changes in gene expression in root primordia of wild type and *coch* mutant lines (Figure 3A). The only interaction of *COCH* with rhizobial infection was the reduced cytokinin response associated with *coch* infections compared to wild type (Figure 6). The hyperinfection phenotype displayed by gibberellin-deficient *na* mutants is known to be associated with ectopic cytokinin response (Velandia *et al*., 2024), so the observed suppression of cytokinin response in *coch* mutants might be why *na coch* double mutants displayed attenuation of infection thread formation and branching (Figure 7A). In addition to playing a key role in nodule organogenesis, *COCH* also promotes bacterial occupancy of the nodules as the infection and fixation zones of *coch* nodules contain less bacteria (e.g. Figure 1E and 5K) and *coch* mutants fix less nitrogen (Liu *et al*., 2023). This may be due to upregulation of defence-related genes (Figure S2B) and mis-regulation of genes with key roles in symbiosome formation such as *DNF1*, *SYT* and *VAMP72* (Gavrin et al., 2017; Ivanov et al., 2012; Van de Velde et al., 2010; Wang et al., 2010)(Figure S2D).

Interestingly, both loss-of-function mutations and overexpression of *COCH* throughout the root lead to the formation of root-like structures emerging from nodules (Figure 4), indicating that *COCH* function is dosage-sensitive and must be tightly regulated during nodule development. This dual phenotype suggests that *COCH* is not merely required to initiate nodules or suppress root identity but rather plays a central role in stabilizing and maintaining the nodule developmental program. Overexpression of *COCH* may perturb downstream regulatory networks or disrupt the balance between auxin and cytokinin signalling pathways, resulting in aberrant organ identity similar to that observed in loss-of-function mutants. Such developmental alterations caused by ectopic expression of homeotic proteins have previously been demonstrated for *KNOX*, *WUSCHEL*, and *PLETHORA* (Aida et al., 2004; Chuck et al., 1996; Gallois et al., 2004).

The negative regulation of nodule number via AON pathway is thought to be triggered by formation of mature nodules (Kassaw *et al*., 2015; Li *et al*., 2009) and has been suggested to act via upregulation of *TML* transcriptional regulator (reviewed by Scott et al., 2024). The fact that *coch* mutants do not produce more nodules than wild type and that there was no additive phenotype in *coch* AON double mutants (Figure 2) indicates the root-nodule hybrids of *coch* mutants can still activate the AON pathway to control nodule number. The failure to induce the expression of *TML2* in mature *coch* nodules (Figure S2D) did not correlate with an increase in nodule number in *coch* mutants and this is consistent with evidence in *Medicago* showing that regulators in addition to TML control nodule number (Schnabel et al., 2023). Interestingly however, we did observe that *coch Psnark* and *coch clv2* double mutants generated a higher proportion of abnormal nodules than *coch* single mutant background. In pea, *Ps*NARK (CLV1) and *Ps*CLV2 putative CLE receptors play a role in regulating shoot meristem proliferation (Scott *et al*., 2024) but these genes did not have an obvious effect on root development (Wang *et al*., 2020). However, in other legumes CLV1 orthologs have been shown to suppress lateral root development (Goh *et al*., 2018; Lagunas *et al*., 2019; Wopereis *et al*., 2000). The additive root-nodule phenotype of *coch Psnark* and *coch clv2* double mutants indicates *Ps*NARK and *Ps*CLV2 may interact with the root program by suppress root identity in developing nodules. Indeed, the upregulation of *KLV1* and *KLV2* expression in developing *coch* nodules (Figure S2D) suggests other potential cross overs between AON pathway and COCH. KLV is a shoot acting receptor that suppresses nodule number and also promotes lateral root formation in *Lotus* (Miyazawa et al., 2010; Oka-Kira et al., 2005) and the KLV ortholog in *Arabidopsis TOAD2/RPK1* suppresses root meristem proliferation (Racolta et al., 2018). Thus, upregulation of *PsKLV1* and/or *PsKLV2* in *coch* nodules may enhance the formation of root-like tissue. These observations point to a broader regulatory role for CLE peptide receptors in organ identity and morphogenesis in root tissues, possibly by acting downstream of COCH. It would be interesting to explore if similar interactions between COCH and CLE peptide signalling occur in the shoot, as both pathways also control shoot organ identity.

Taken together, our findings provide direct evidence that COCH, modulates cytokinin and auxin pathways at multiple levels—transcriptional regulation, hormone levels, and spatial hormone response—to direct nodule formation and preserve nodule identity (Figure 8). In our studies RNAseq and hormone analysis were analysed at organ level and hormone response was spatially delineated, in the future it would be enlightening to monitor spatial levels of transcription and hormone level to resolve the precise mode of action of *COCH* to regulate hormone dynamics during nodule development. Members of the NBCL gene family, including COCH, function as transcriptional co-regulators that influence the development of diverse shoot organs—such as leaves, flowers, and inflorescences—across monocots and dicots, both legumes and non-legumes (Couzigou *et al*., 2012; Khan et al., 2014; Liu *et al*., 2023; Magne *et al*., 2018a) and it would be interesting to explore if COCH regulates common targets in auxin and cytokinin pathways to shape shoot development.

## Methods

### Plant material, germplasm generation and growth conditions

The *Pisum sativum L.* lines used in this study were *coch-JI216* and *coch-JI2757* and their respective wild type lines cv. Weitor and Parvus (Blixt, 1967; Couzigou *et al*., 2012). The gibberellin-deficient mutant *na-1* derived from WT line WL1769 (Davidson et al., 2003), *Psnark* (P88, formerly *sym29*, disrupted in *PsCLV1*) and *Psclv2* (P64, formerly *sym28*) derived from the parental line ‘Frisson’ (Duc and Messager, 1989; Krusell et al., 2002; Krusell et al., 2011; Sagan and Duc, 1996; Schnabel et al., 2011). The double mutant line *na coch* were generated from cross between *coch-JI2165* and *na-1*. The double mutant lines *Psnark coch* and *Psclv2 coch* were generated from cross between *coch-JI2165* and *Psnark* or *Psclv2*. Double mutant, single mutants and wild type derivatives from these crosses were selected in the F2 generations using reduced stipules phenotype to select *coch,* dwarfism for *na* and RFLP markers to select *Psnark* or *Psclv2*. Primers for PCR amplification of template are listed in Table S1. Restriction digests were carried out using BsmAI for *PsNARK* and BtsCI for *PsCLV2*. Unless otherwise stated plants were grown in glasshouse conditions as described by Foo and Davies (2011) in sterile conditions and inoculated at day 7 with *Rhizobium leguminosarum* bv *viciae* strain RLV248 containing the plasmid Phc60 for GFP constitutive expression (RLV248G) (Cheng and Walker, 1998) as described by Velandia *et al*. (2024) and received modified Long Ashton nutrients with no nitrogen weekly (Hewitt, 1966). To characterise root architecture of *coch* lines, plants were grown with respective wild type progenitors in sterile conditions for 28 days and received sterile Long Ashton nutrients with no nitrogen 14 and 20 days after planting. For AM study, plants were inoculated with *Rhizophagus irregularis* and grown with modified Long Ashton nutrients with 10 mM KNO3 and 0.05 mM NaH_2_PO_4_ weekly as outlined in Foo et al. (2013).

### Examination of nodulation structures and root architecture

For nodulation experiments in AON *coch* double mutants, plants were harvested 3 weeks after rhizobial inoculation and nodules were scored by eye and classified as mature nodule (typical mature pea nodule) or root-nodule hybrid (with protruding root-like structures characteristic of coch mutant; Ferguson and Reid, 2005). For all other experiments roots were examined using a Zeiss Axioscope5 fluorescence microscope. The total number of IT, developing nodules (not yet emerged from the root profile), mature nodules and abnormal nodules (including root-nodule hybrids) were counted per cm of root length. For *na coch* mutants, branched IT were also counted and due to the reduced root length of plants carrying the *na* mutation structures were expressed the number of structures per cell as outlined in Velandia *et al*. (2024). For *TCSn::GUS* transformation experiments the presence of GUS staining associated with each structure was also recorded. Root architecture was examined in Table S2 was performed measuring the length of the top 6 secondary roots, the number of tertiary roots on the longest secondary root and the length of the longest tertiary root on the longest secondary root.

### Construct assembly and hairy root transformation

Specific promoters and genes where inserted in the Pcambia_CR1 (PCR1) vector (Sevin-Pujol et al., 2017) using Golden Gate system, following the instructions of Golden Gate assembly kit (New England Biolabs). The *PsCOCH* coding region (Psat3g077680) (1467bp) was obtained from pea root cDNA. The *AtCASP1* endodermal promoter (Roppolo et al., 2011) and *CO2* promoter, were obtained directly from a PBS plasmid kindly provided by Sandra Bensmihen (Sevin-Pujol *et al*., 2017). The two component signalling (*TCSn*) cytokinin responsive promoter was supplied by Bruno Muller (Zürcher et al., 2013) and was linked to the *GUS* reporter in the PCR1 vector. The *DR5* promoter was obtained from the pGreenIIM DR5v2-ntdTomato/DR5-n3GFP (Liao et al., 2015) and fused with *GUS* reporter gene in the PCR1 vector. PCR fragments for *TCSn, DR5* and *35S* promoter regions and *GUS* and *COCH* genes where generated using primers containing the BsaI recognition site and matching overhangs (Table S1). Pea roots were transformed with *Agrobacterium rhizogenes* (ARqua1) carrying the corresponding construct according to Velandia *et al*. (2024). Plants were harvested three weeks post rhizobial inoculation. For harvesting, individual roots regenerated from the callus of 10– 20 plants were screened for transformation using the DsRED fluorescent marker. Transformed roots were then fixed in 1% paraformaldehyde and stored at 4 °C for subsequent scoring. For *TCSn* and *DR5* studies, roots were stained for GUS before fixation in paraformaldehyde as described in Velandia *et al*. (2024).

### Root sectioning

Roots were removed from paraformaldehyde, trimmed and placed into 2.5M sucrose overnight. Tissue was embedded in 1:1 mixture of 2.5M sucrose:OCT (Scigen) in a cryostat mold, snap frozen and 16μm sections were made using Epredia Cryostar HM525 NX (Epredia, UK). Sections were briefly rinsed in water and mounted on slides.

### RNA seq studies

Wild type (Weitor) and *coch JI-2165* plants were inoculated with RLV248G 14 days after planting and grown under glasshouse conditions as described. When plants were 6 weeks old, roots were washed and for each genotype three tissue types were harvested; root tips, developing nodules (< 2mm in size, with a small amount of mature root attached) and mature nodules (> 2mm in size, with a small amount of mature root attached). In *coch* plants, mature nodules consisted of root–nodule hybrids along with a few visually normal nodules. For each tissue type and genotype combination, four biological replicates were collected, each comprising tissue from 2–3 plants. Additional replicate samples of mature nodules were collected separately for hormone analysis, as described below. Tissue was frozen in liquid-N and RNA was extracted using ISOLATE II RNA mini kit (Bioline). RNA samples were processed for mRNA library preparation (poly(A) enrichment) and sequenced on a NovaSeq X Plus platform (PE150) by NovogeneAIT (Singapore). Quality control assessment of the RNA-seq data was conducted using the FastQC tool (Andrews, 2017). Based on the quality control analysis, reads were subsequently processed using Trimomatic (Bolger and Giorgi, 2014). The sequences underwent pseudoalignment against the pea (*Pisum sativum*) Cameor transcriptome (Alves-Carvalho et al., 2015) using Kallisto (Bray et al., 2016). The differential expression analysis was done using Sleuth package (Pimentel et al., 2017) and the differentially expressed genes were obtained by pairwise comparison of each treatment against the wild type root primordia treatment as a control. The selection of DEGs was based on the false discovery rate corrected p-values under 0.05 and log_2_ fold of change over 1. GO enrichment analysis was performed using TopGO package (Alexa et al., 2006).

### Hormone analysis

Auxin and cytokinin extraction and quantification was carried out as outlined in Bound et al. (2022) with stable isotope-labelled internal standards added to each sample. Samples were then analysed using a Waters Acquity H-Class UPLC instrument coupled to a Waters Xevo triple quadrupole mass spectrometer. For all samples, peak areas for endogenous and labelled hormones were compared and combined with the FW of samples to calculate ng/g FW.

## Author contributions

E.F. conceived the project. N.S., T.S., K.V., A.C., A.M, and E.F. performed the experiments and analysed the data. E.F wrote the manuscript with input from all authors.

## Acknowledgements

We thank Maria Soto (University of Granada) for the gift of GFP-labelled *Rhizobium*, Sandra Bensmihen (INRA, France) for the *pCAMBIA CASP* and *CO2* construct, Bruno Mueller (Zurich-Basel Plant Science Center, Department of Plant and Microbial Biology, UZH) for the *TCSn* promoter, and David Nichols (CSL, UTAS) for expert assistance with hormone analysis. We thank Tracey Winterbottom and Sarah Kane for assistance with plant husbandry.

## Supporting Information

**Supplementary Table 1.**
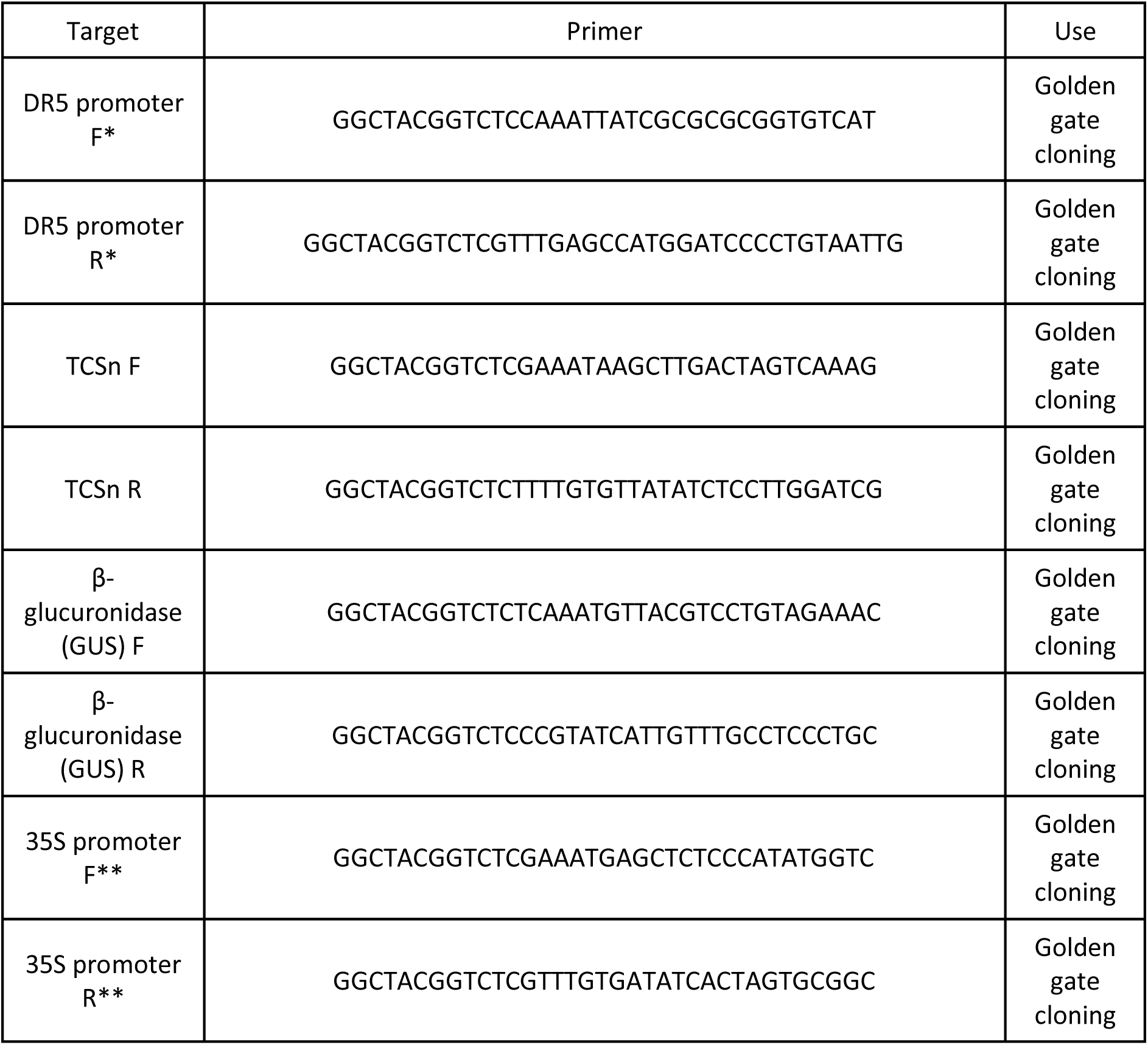
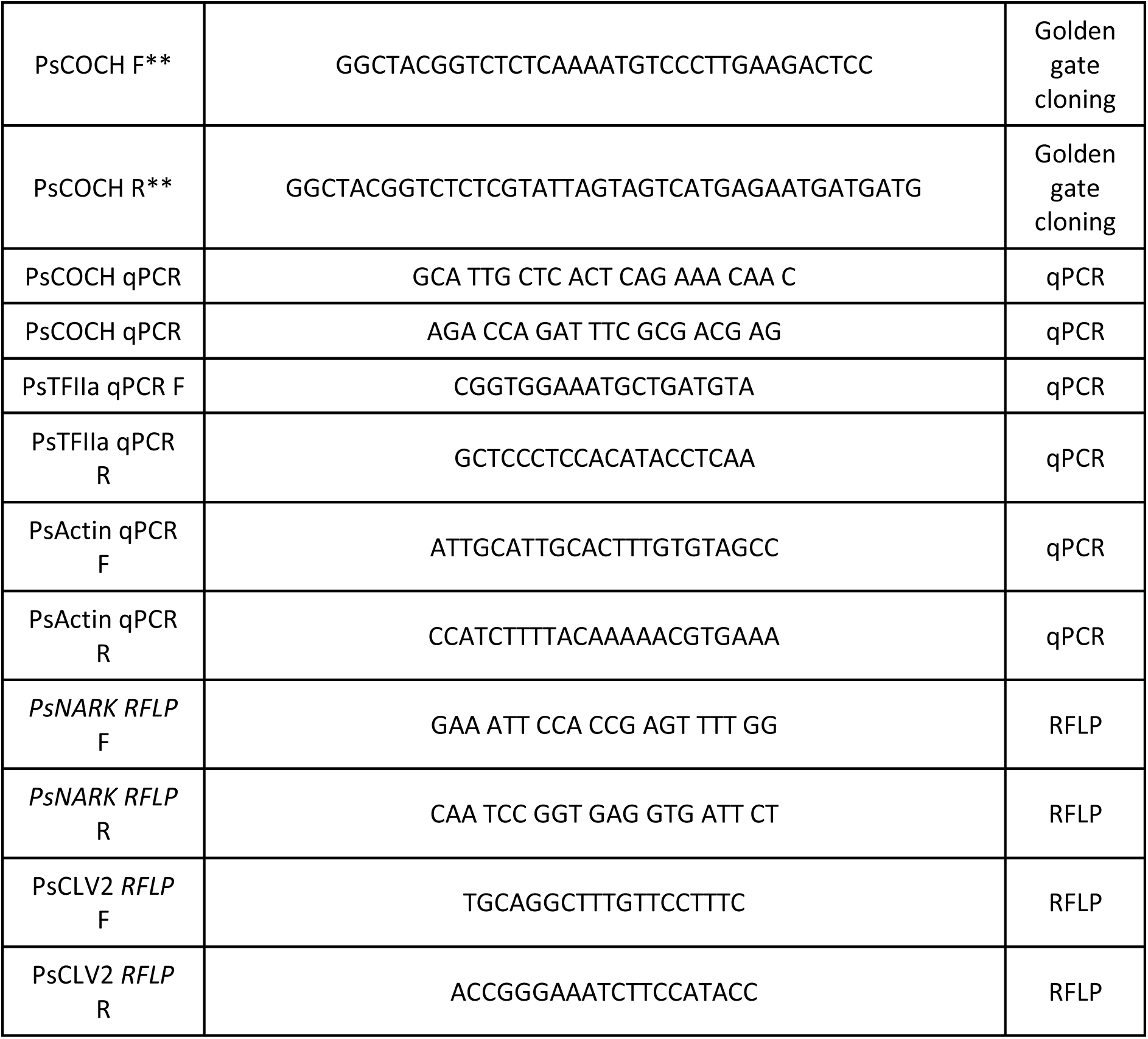
List of primers used for Golden Gate assembly, qPCR, and RFLP analysis. This table includes the sequences and relevant notes for primers used in Golden Gate cloning, quantitative Polymerase Chain Reaction (qPCR), and restriction fragment length polymorphism (RFLP) analysis. Primer sequences are listed in the 5′ to 3′ direction. *Due to the highly repetitive nature of the pGreenIIM DR5v2-ntdTomato/DR5-n3GFP construct, an internal region containing the promoter was first amplified using flanking primers designed outside the repetitive sequence. This PCR product was then used as a template for amplification with Golden Gate-compatible primers for subsequent cloning: DR5-flank-F : ACCTGTTCCTGGGGCATG, DR5-flank-R : CTTGCTCACCATGGATCTCT. ** Fragments were domesticated prior to cloning to remove an internal BsaI restriction site.

**Supplementary Table 2.** Root phenotypes of *coch*-JI2165 and *coch*-JI2757 mutants and their corresponding isogenic wild-type lines, Weitor and Parvus. Plants were grown under sterile conditions for 28 days. Values are presented as mean ± standard error (SE), *n* = 10–16. Asterisks indicate statistically significant differences from the respective wild-type line, based on Student’s *t*-test (\**p* < 0.05; \*\**p* < 0.01). The length of secondary roots corresponds to the average length of the six longest secondary roots. The number of tertiary roots was counted on the longest secondary root. The length of the tertiary root refers to the longest tertiary root on that same secondary root.

| Genotype | Tap root length (cm) | Length secondary root (cm) | No. secondary roots/cm tap root | No. tertiary roots/cm secondary root | Length tertiary root (cm) | Shoot DW (mg) | Root DW (mg) | Shoot:root ratio |
| --- | --- | --- | --- | --- | --- | --- | --- | --- |
| Weitor | 20.3 ± 1.5 | 18.1 ± 0.7 | 2.8 ± 0.3 | 1.9 ± 0.1 | 4.9 ± 0.7 | 0.2 ± 0.01 | 0.1 ± 0.003 | 2.9 ± 0.2 |
| *coch-JI2165* | 21.8 ± 1.2 | 17.0 ± 1.3 | 3.9 ± 0.2** | 1.8 ± 0.1 | 4.7 ± 0.5 | 0.2 ± 0.01 | 0.07 ± 0.006 | 3.1 ± 0.2 |
| Parvus | 27.5 ± 2.4 | 14.1 ± 0.9 | 3.3 ± 0.2 | 1.8 ± 0.1 | 5.4 ± 0.8 | 0.2 ± 0.02 | 0.09 ± 0.01 | 2.1 ± 0.1 |
| *coch-JI2757* | 21.0 ± 1.2* | 16.9 ± 0.8* | 3.5 ± 0.2 | 1.9 ± 0.1 | 7.2 ± 0.6 | 0.3 ± 0.02* | 0.11 ± 0.1 | 2.3 ± 0.9 |

**Supplemental Table 3.**
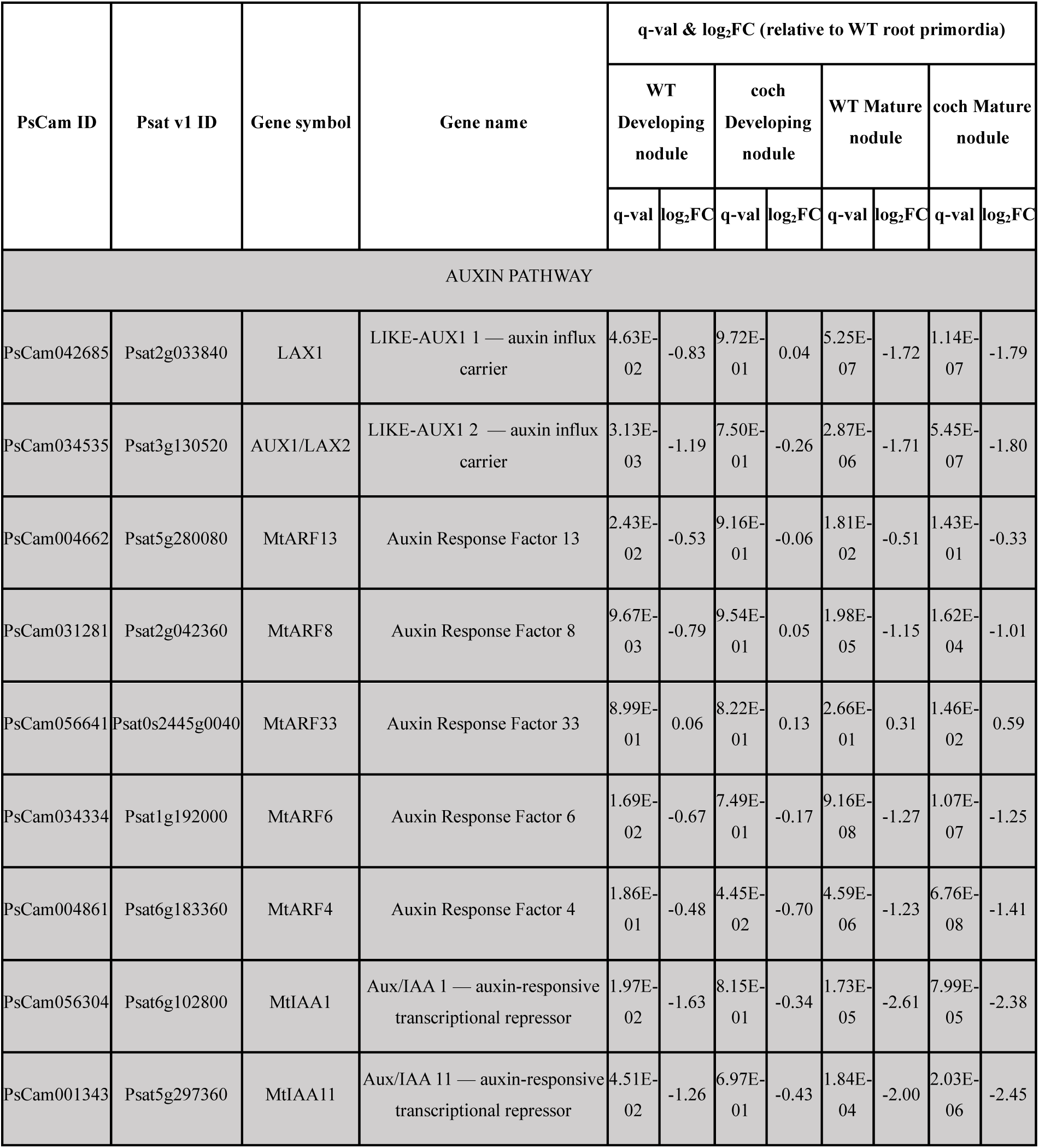
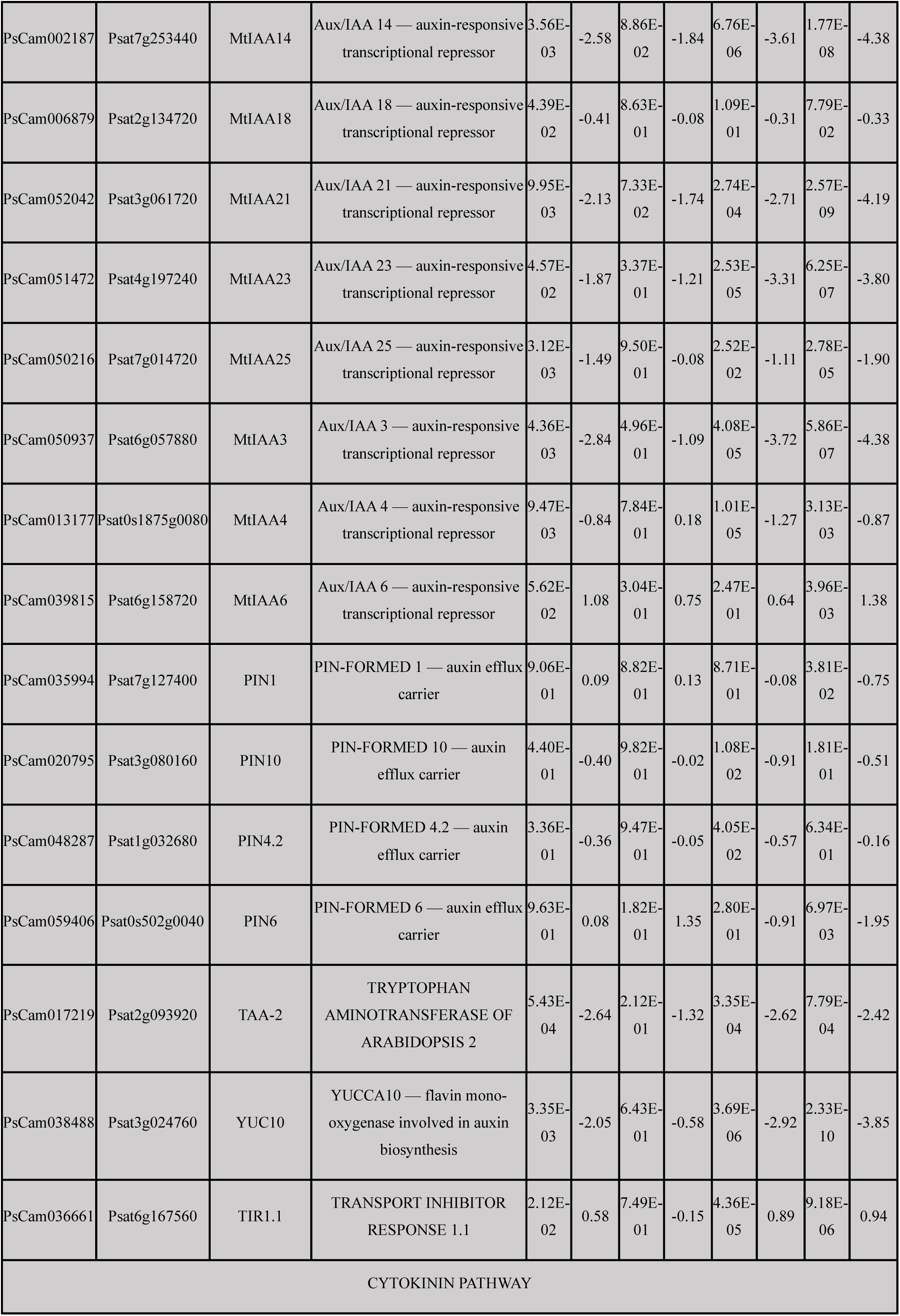
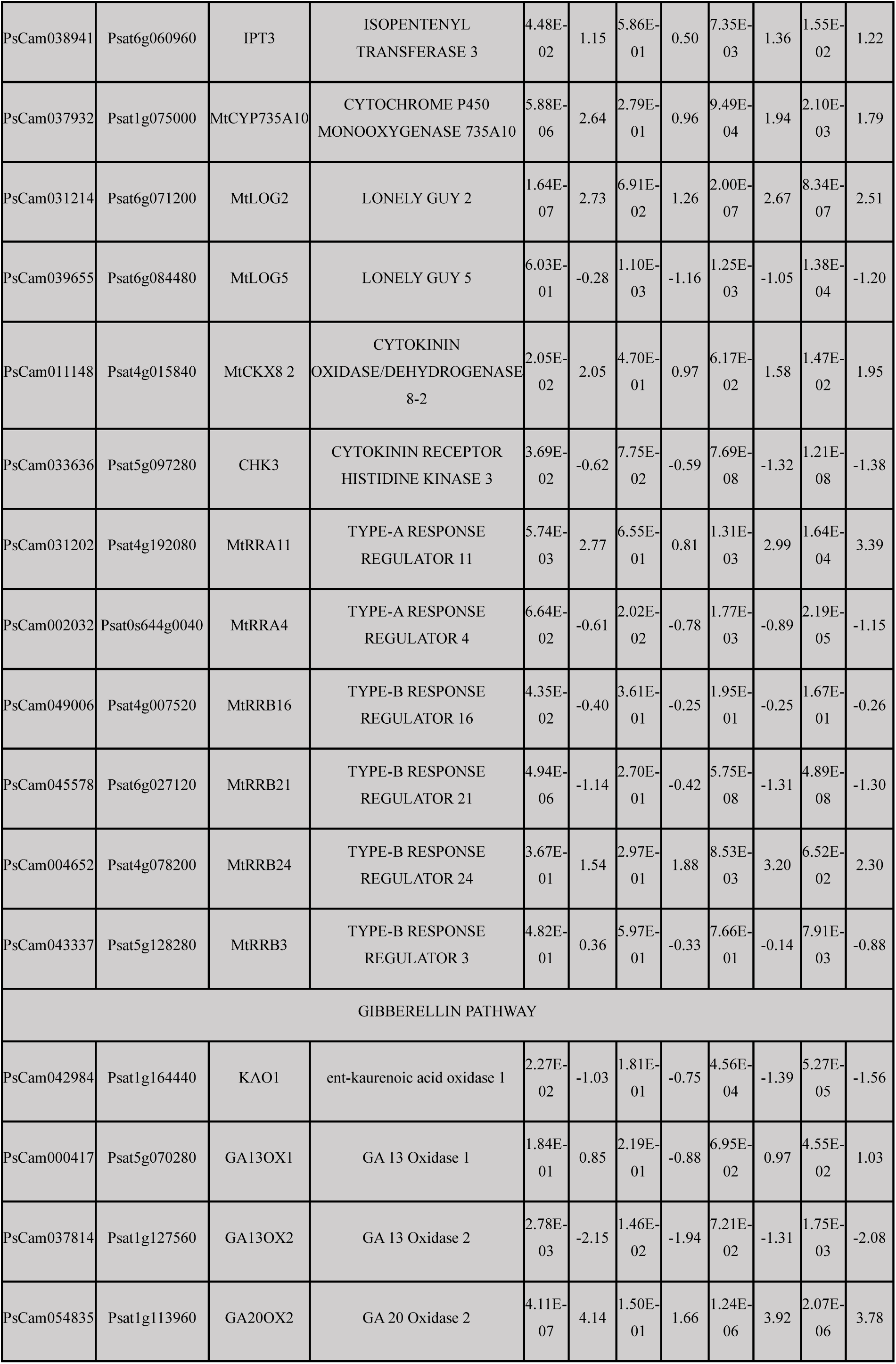
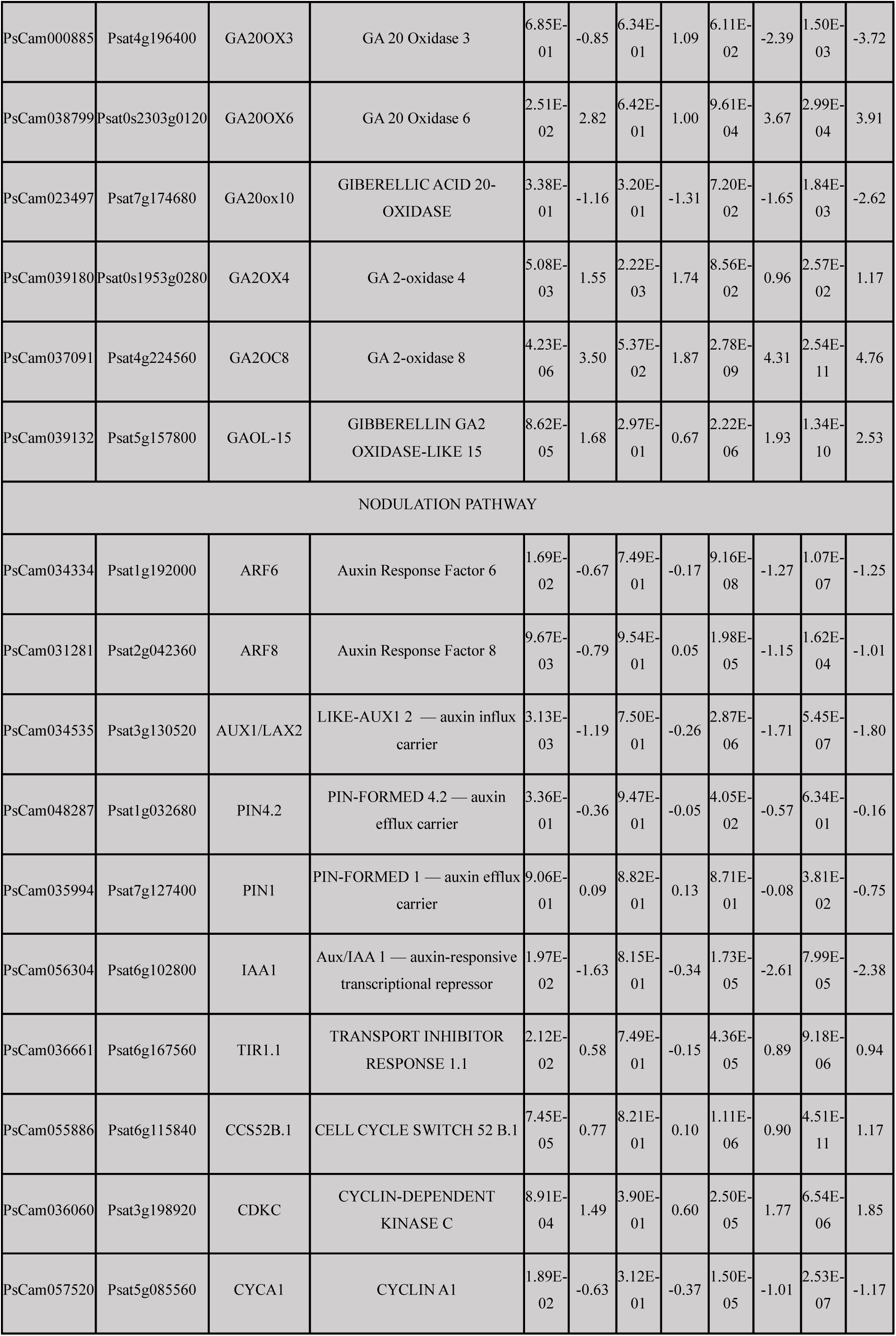
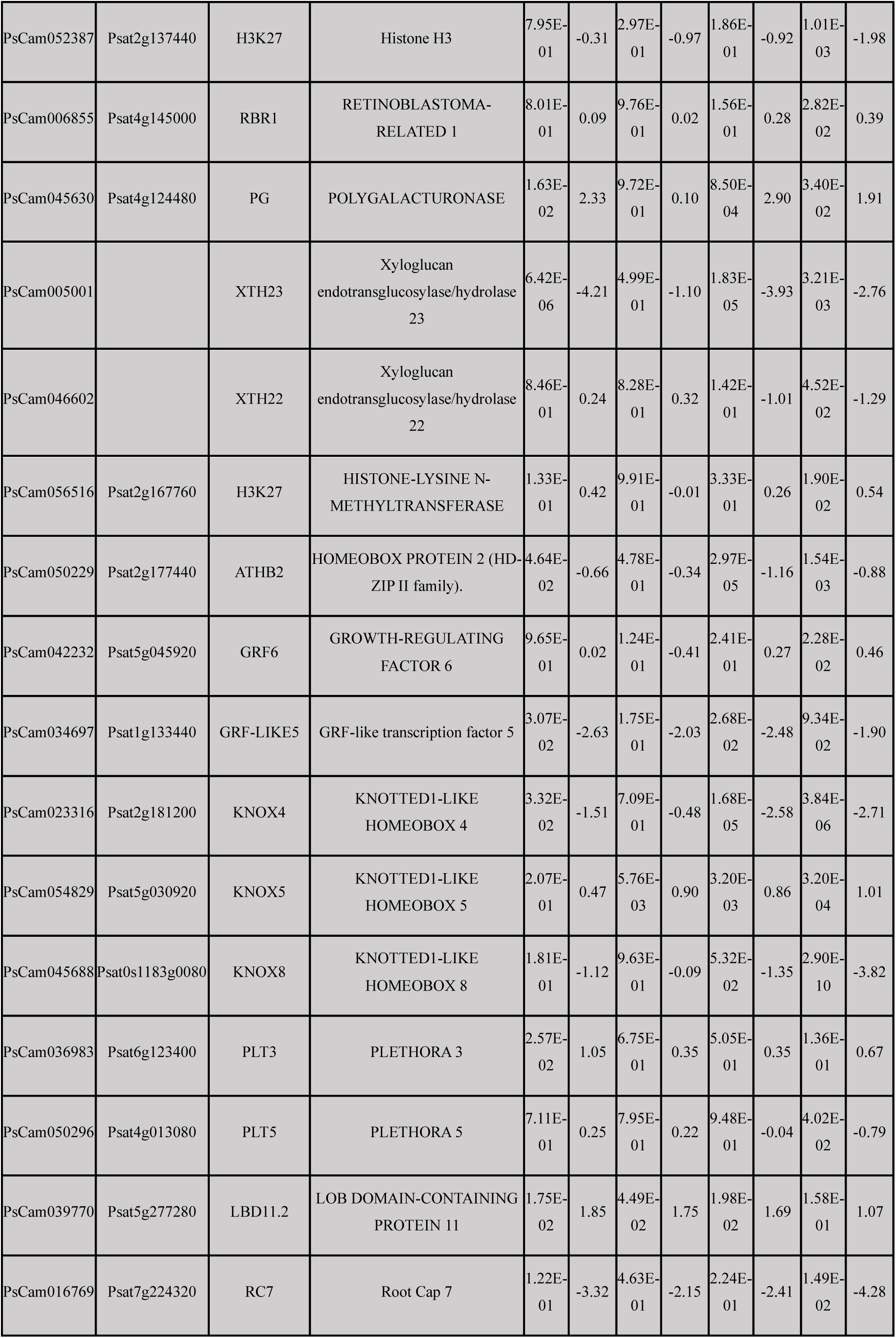
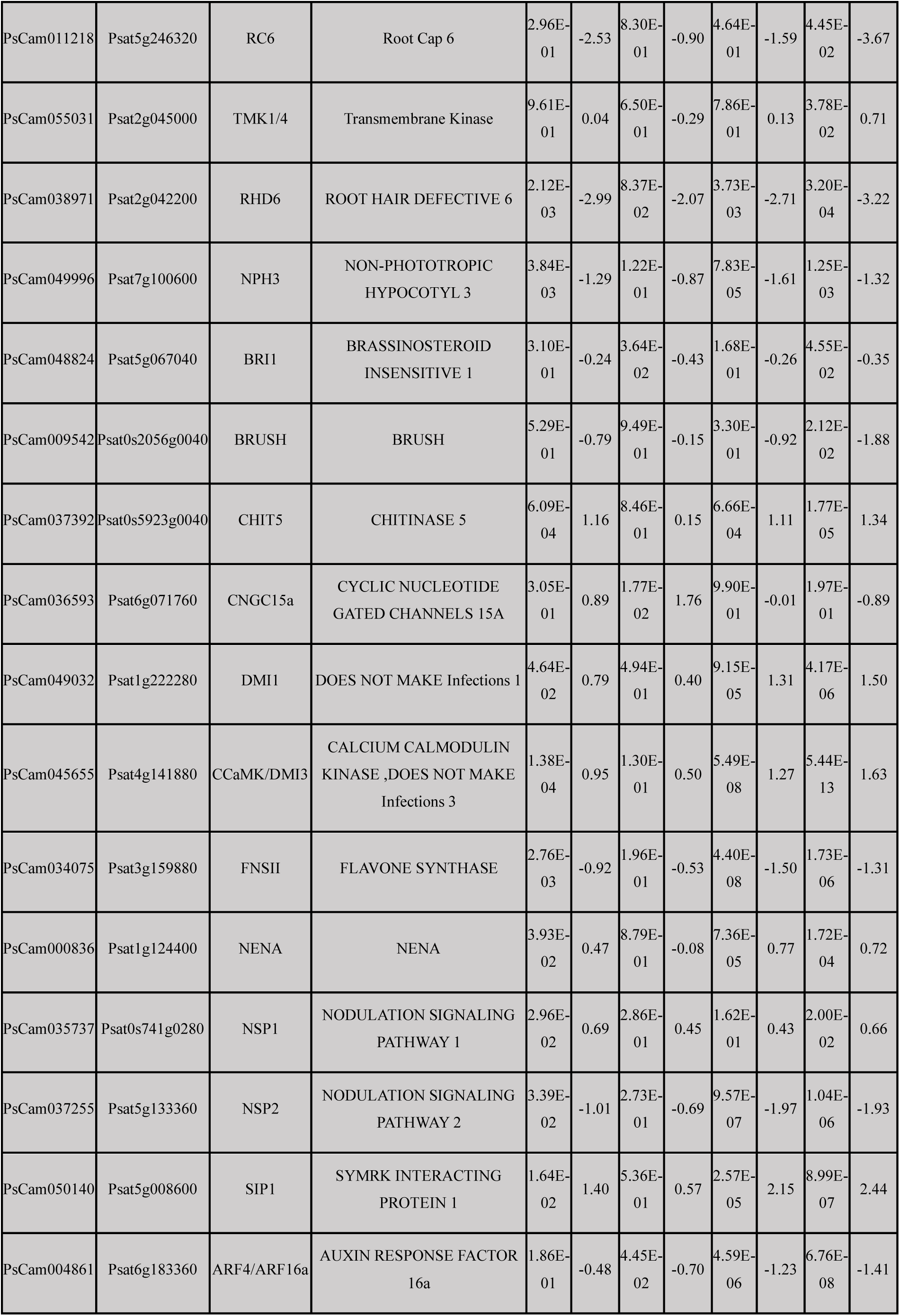
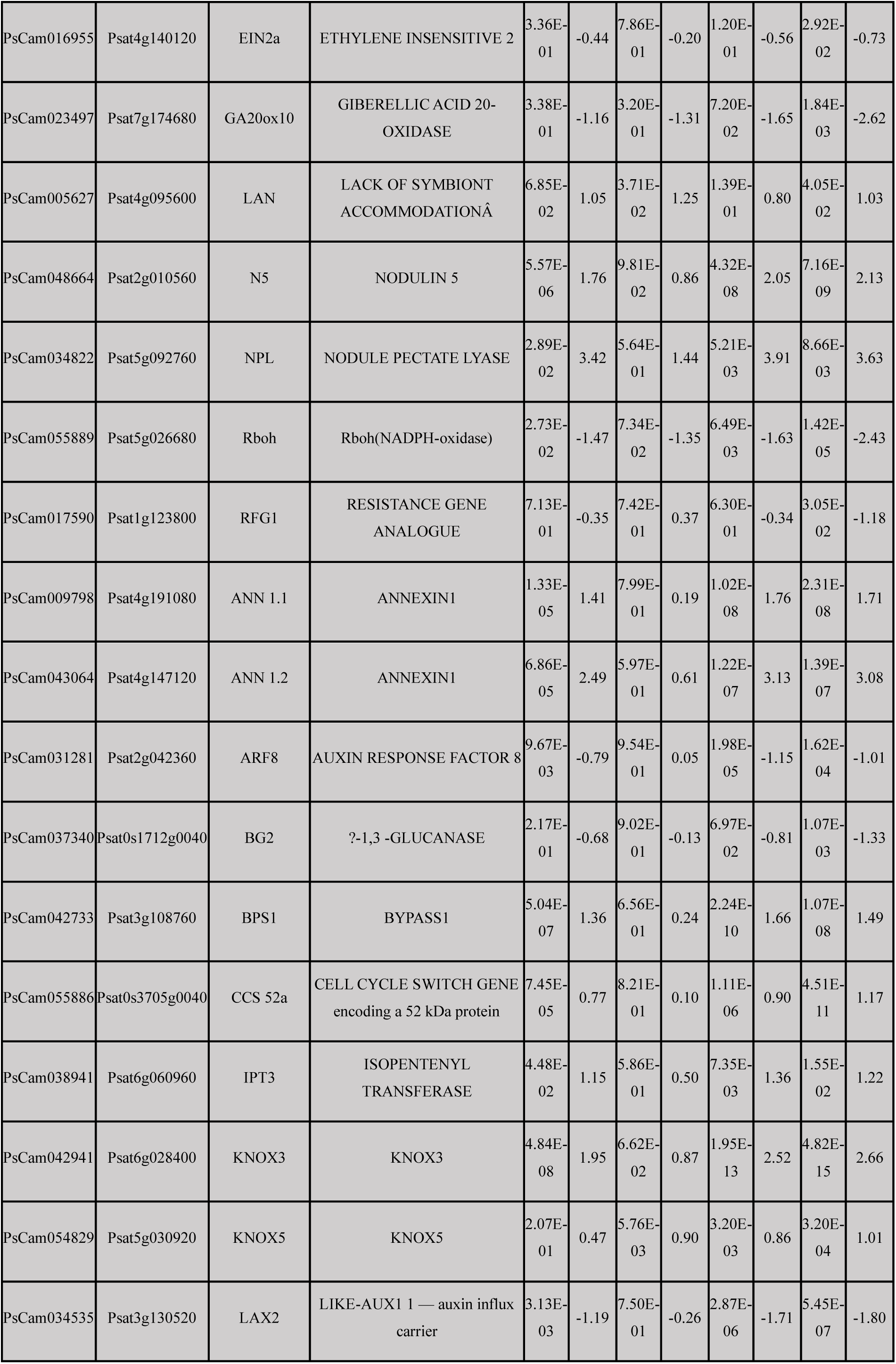
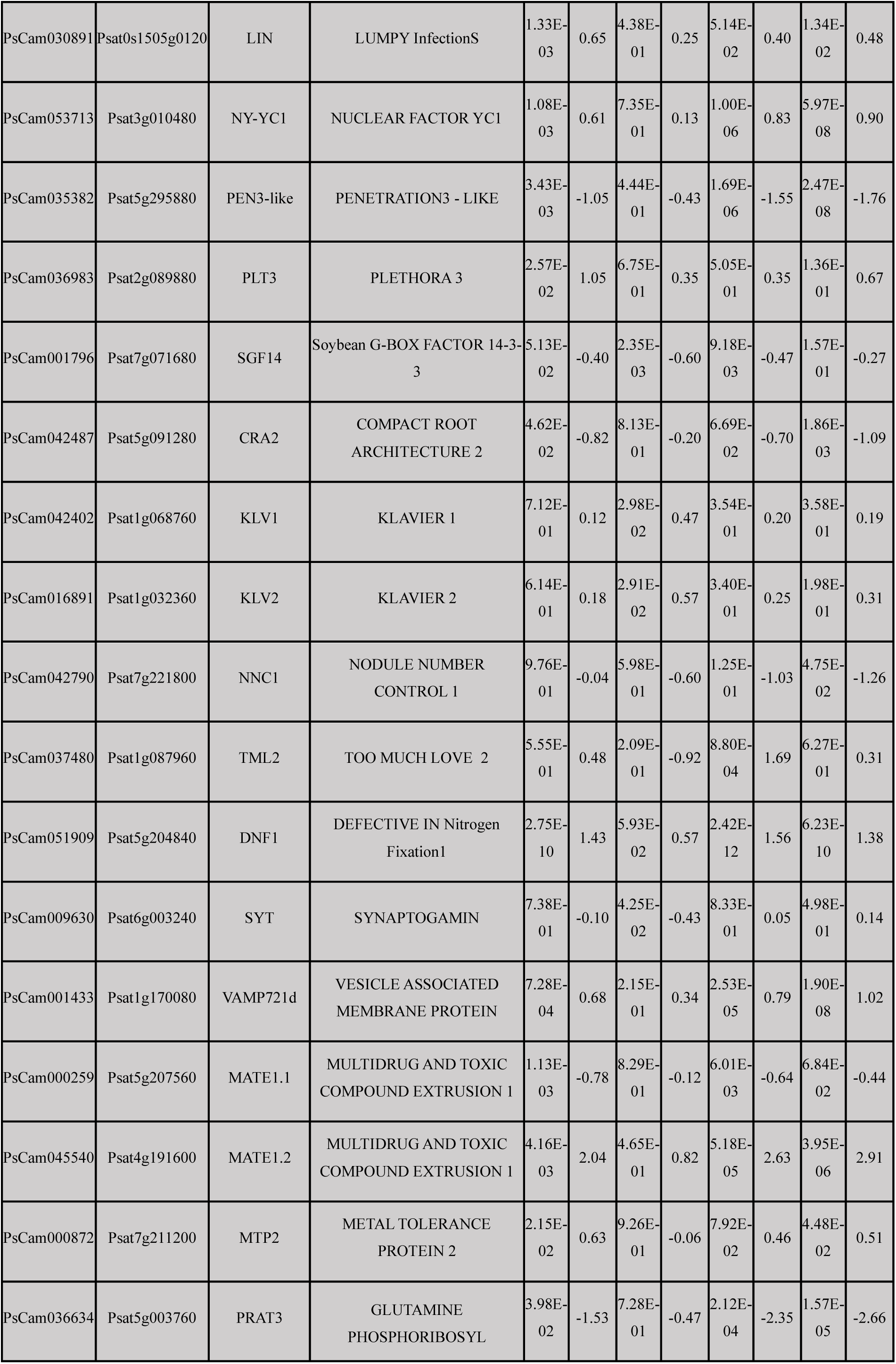
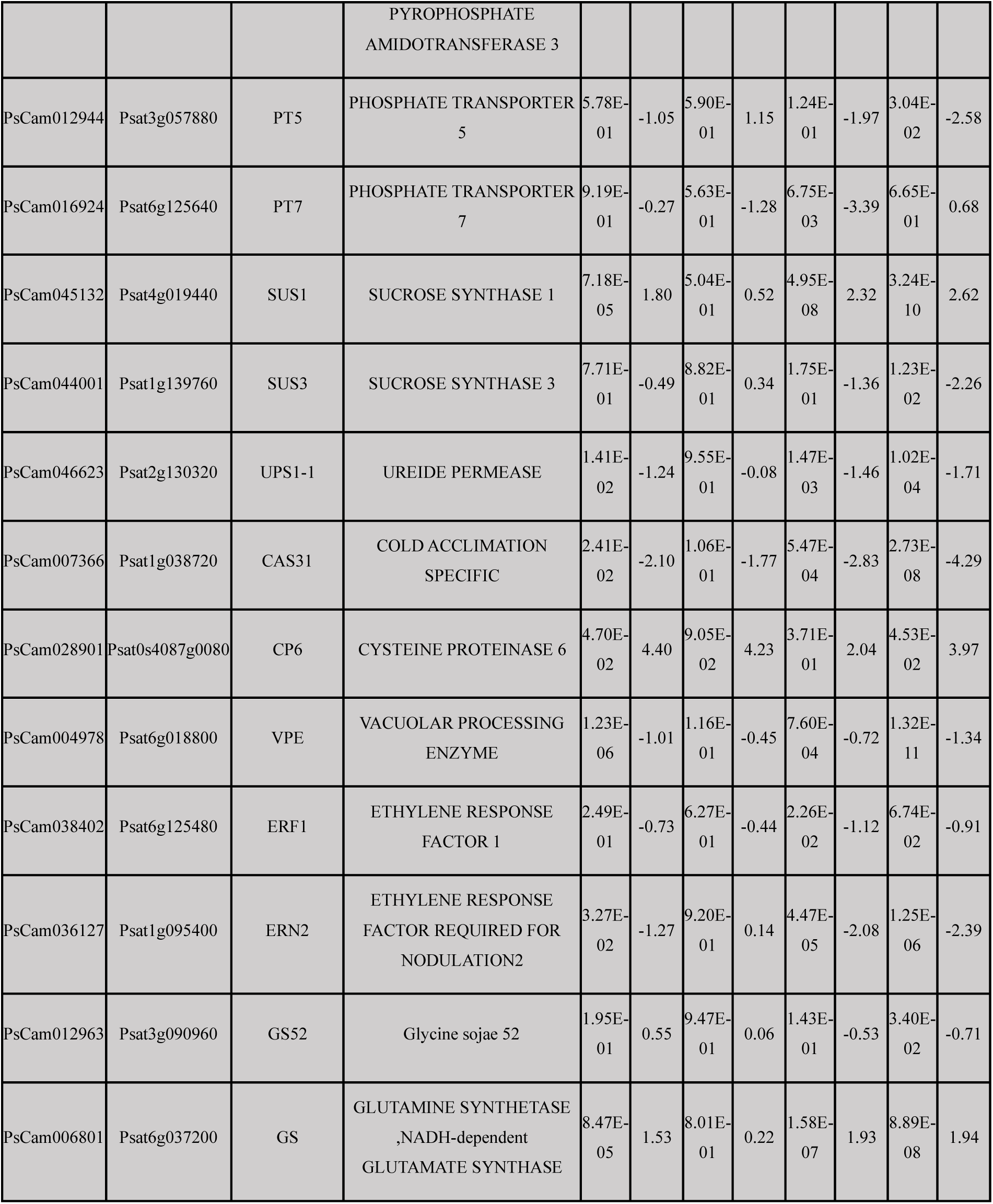
Genotype-specific DEGs across developmental pathways in developing and mature nodules (vs WT root primordia) Pisum sativum genes that are differentially expressed only in WT or only in *coch* (genotype-specific) when comparing developing nodules and mature nodules to WT root primordia. Genes that were differentially expressed in both genotypes within the same tissue, or did not differ between WT and *coch* for that tissue, are excluded. Columns list PsCam transcript ID, Psat v1 gene ID, gene symbol, gene name, and per genotype tissue log2FC and q-value, as defined in Methods.

**Supplementary Figure 1.**
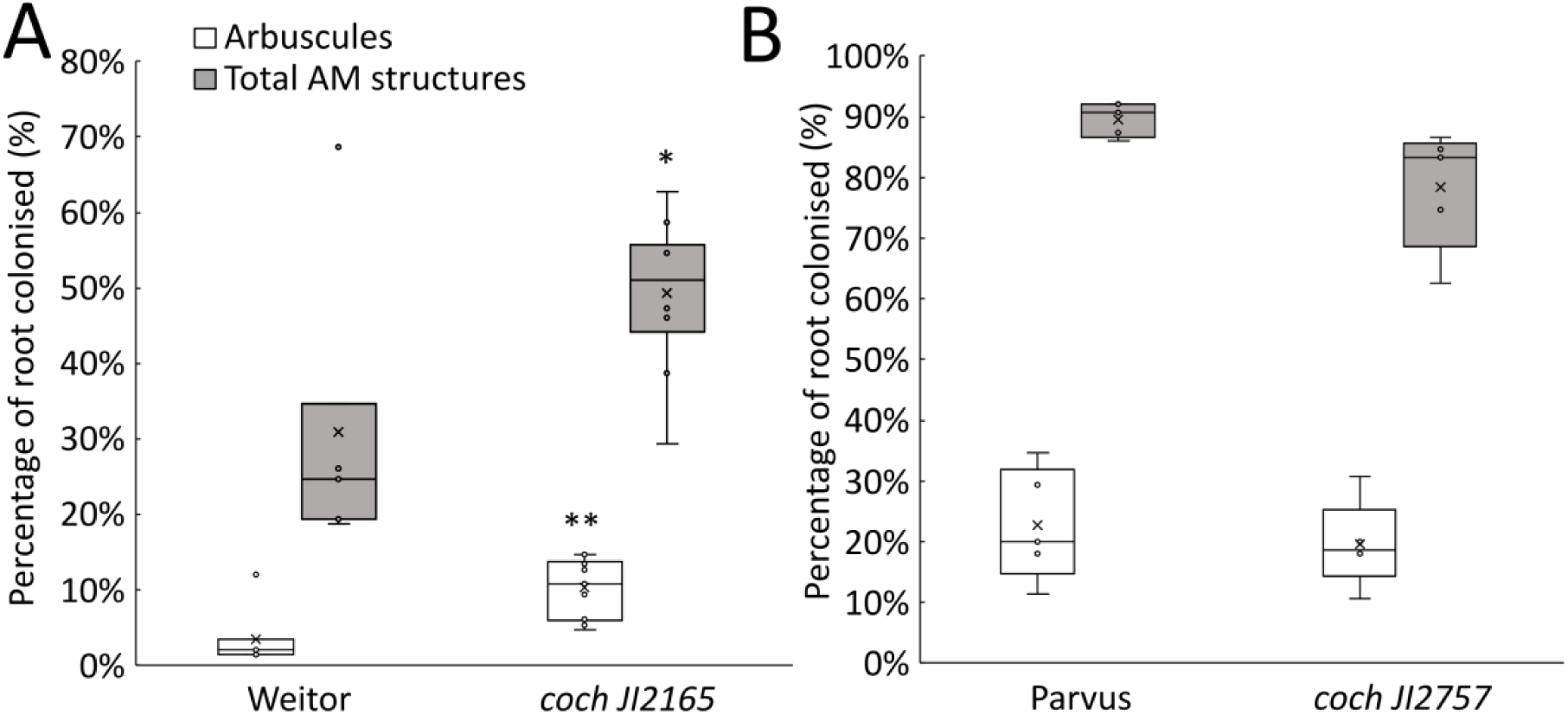
Mycorrhizal phenotypes *coch*-JI2165 and *coch*-JI2757 mutants and their corresponding isogenic wild-type lines, Weitor and Parvus. (A-B) Percentage of roots colonised with arbuscules or total arbuscular mychorrizal (AM) structures (arbuscules, vesicles and hyphae) in *coch-JI2165* and wild type Weitor (A) and *coch-JI2757* and wild type Parvus (B). Data are shown as box plots: boxes represent the interquartile range, horizontal lines show the median, and whiskers indicate the full data range. *n* = 7–10; *p* < 0.05 (*), *p* < 0.01 (**), Student’s *t*-test.

**Supplementary Figure 2.**
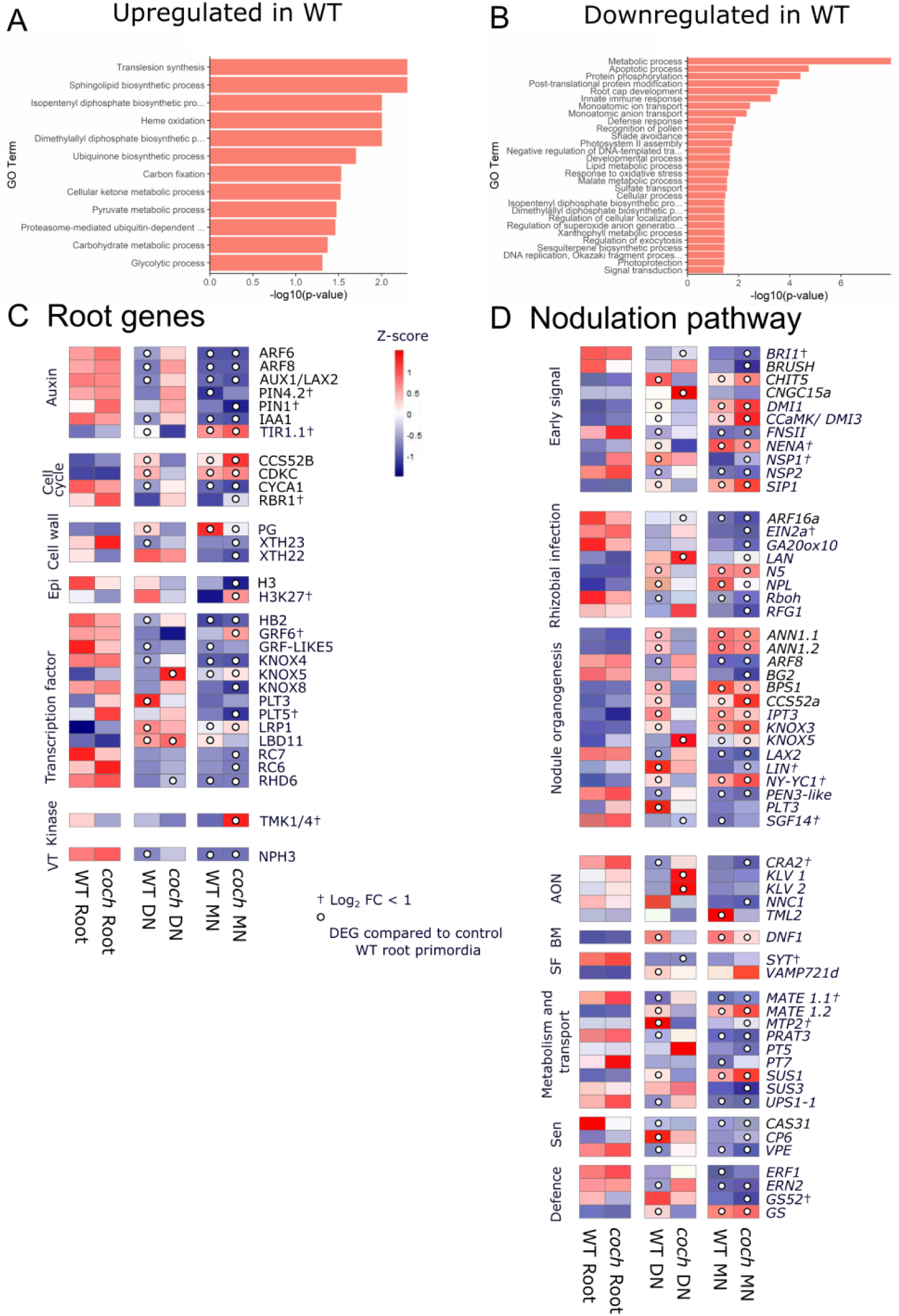
Gene ontology enrichment and expression profiles of root and nodulation-related genes. (A–B) Gene Ontology (GO) enrichment analysis of genes uniquely upregulated (A) or downregulated (B) in wild-type plants in developing and mature nodules, which showed no response in *coch* mutants. (C–D) Heatmaps showing the expression of genes involved in lateral root formation (C) and nodulation (D) that were differentially expressed between wild-type and *coch* mutants in either developing or mature nodules. The heatmap colour scale represents Z-scores of normalized gene expression. White circles within each heatmap cell indicate that the gene in that specific condition was significantly differentially expressed relative to the wild-type root primordia (q-value < 0.05). Genes marked with a cross (†) indicate that although differentially expressed, their absolute log₂ fold change (log₂FC) was < 1. Samples were collected from 6-week-old plants, 4 weeks post-inoculation with *Rhizobium*. Tissues included root tips, developing nodules, and mature nodules for wild-type, and abnormal nodules for *coch* mutants. Abbreviations: Epi – Epigenetics, VT – Vesicular transport, AON – Autoregulation of nodulation, BM – Bacterial maturation, SF – Symbiosome formation, Sen – Senescence. DN: Developing nodule. MN: Mature nodule. Gene symbols and full names are listed in Table S3.

**Supplementary Figure 3.**
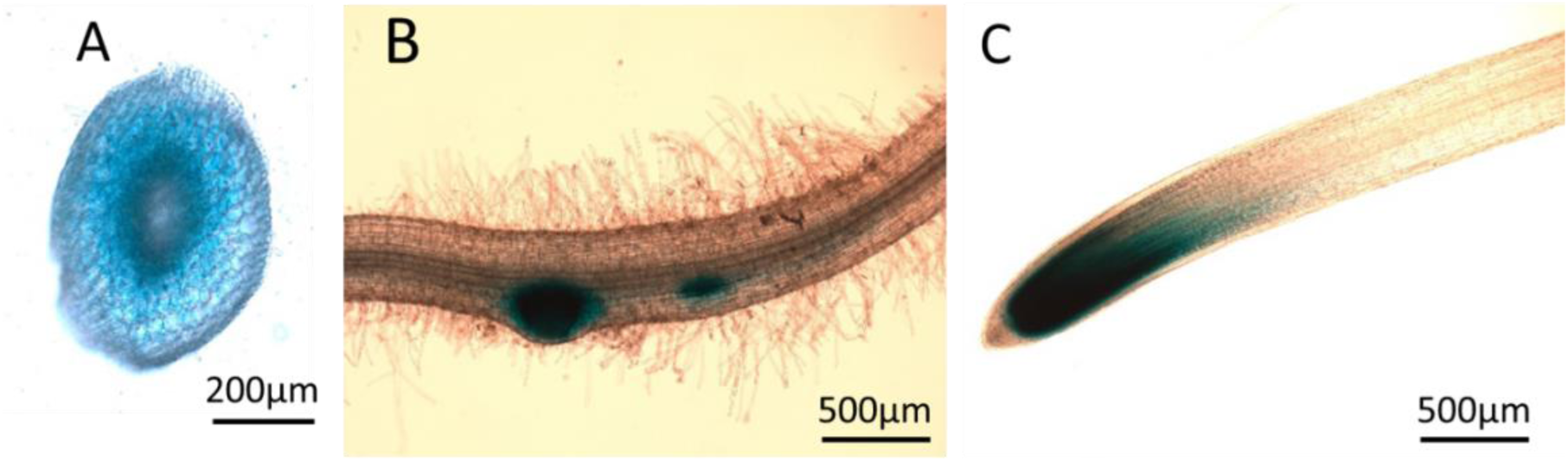
Tissue-specific expression of the *CO2* promoter in pea (*Pisum sativum*) used in Figure 4. Hairy root-transformed *Pisum sativum* (cv. Torsdag) plants were used to analyse *CO2* promoter activity using a *CO2::GUS* construct (β-glucuronidase reporter gene). GUS activity is indicated by blue staining. (A) Cross-section of a root showing *CO2*::*GUS* expression in the cortex at the basal meristematic zone and above. (B) *CO2*::*GUS* expression in early nodule primordia. (C) *CO2*::*GUS* activity in the root tip.

**Supplementary Figure 4.**
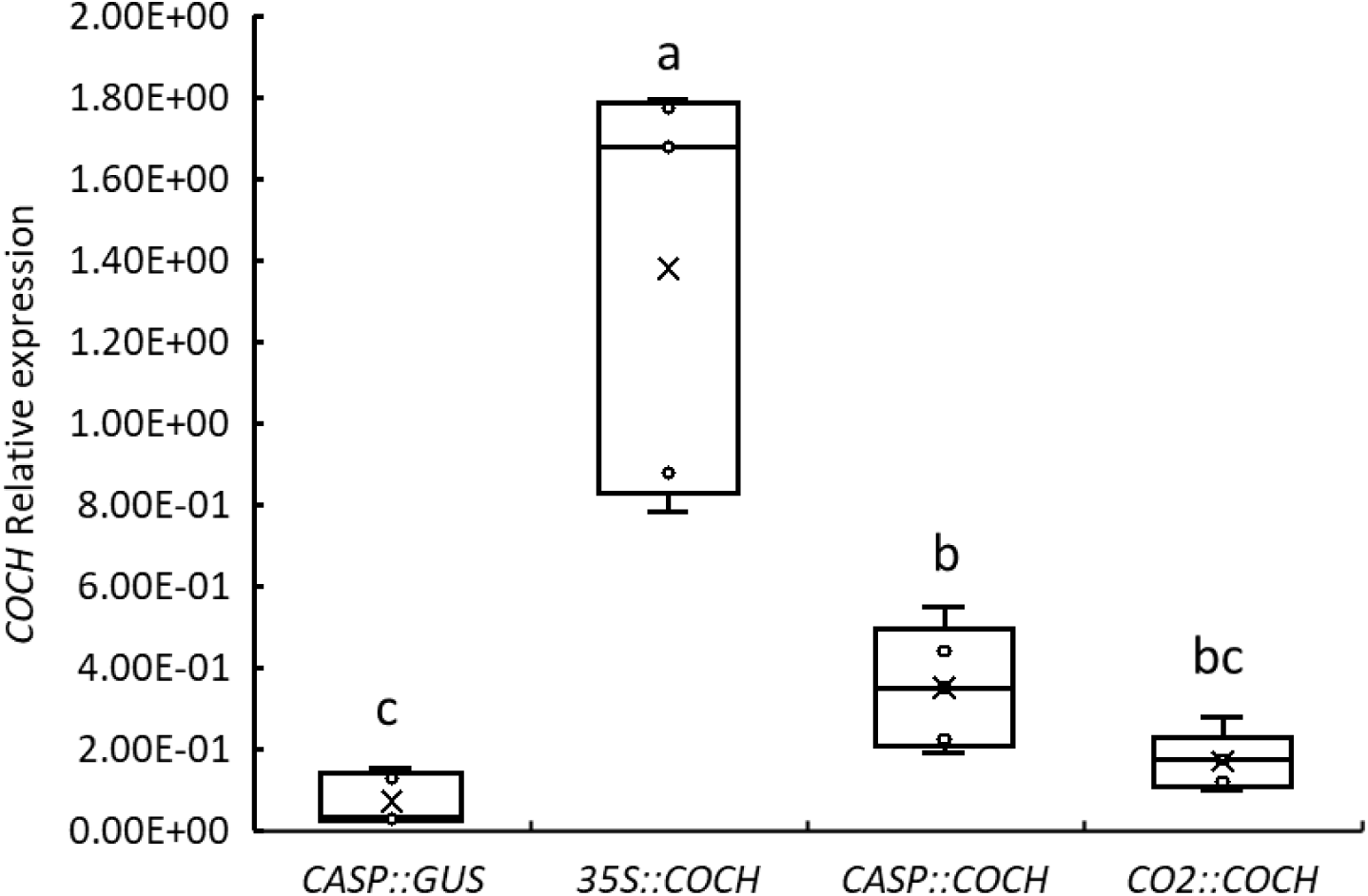
Relative expression of *PsCOCH* in ectopic expression hairy root transformation experiment shown in Figure 4. Relative expression of *PsCOCH* in transformed pea hairy roots expressing different constructs (*CASP::GUS, 35S::COCH, CASP::COCH, CO2::COCH*). Expression was normalized to the geometric mean of two housekeeping genes: *TFIIa* and *Actin*. Whole transformed roots were harvested 3 weeks post-inoculation with *Rhizobium leguminosarum* for RNA extraction and qPCR. Data are shown as box plots: boxes represent the interquartile range, horizontal lines show the median, and whiskers indicate the full data range. *n* = 5. Different letters indicate statistically significant differences between constructs as determined by one-way ANOVA followed by Tukey’s HSD post hoc test (p < 0.05).

**Supplemental Figure 5.**
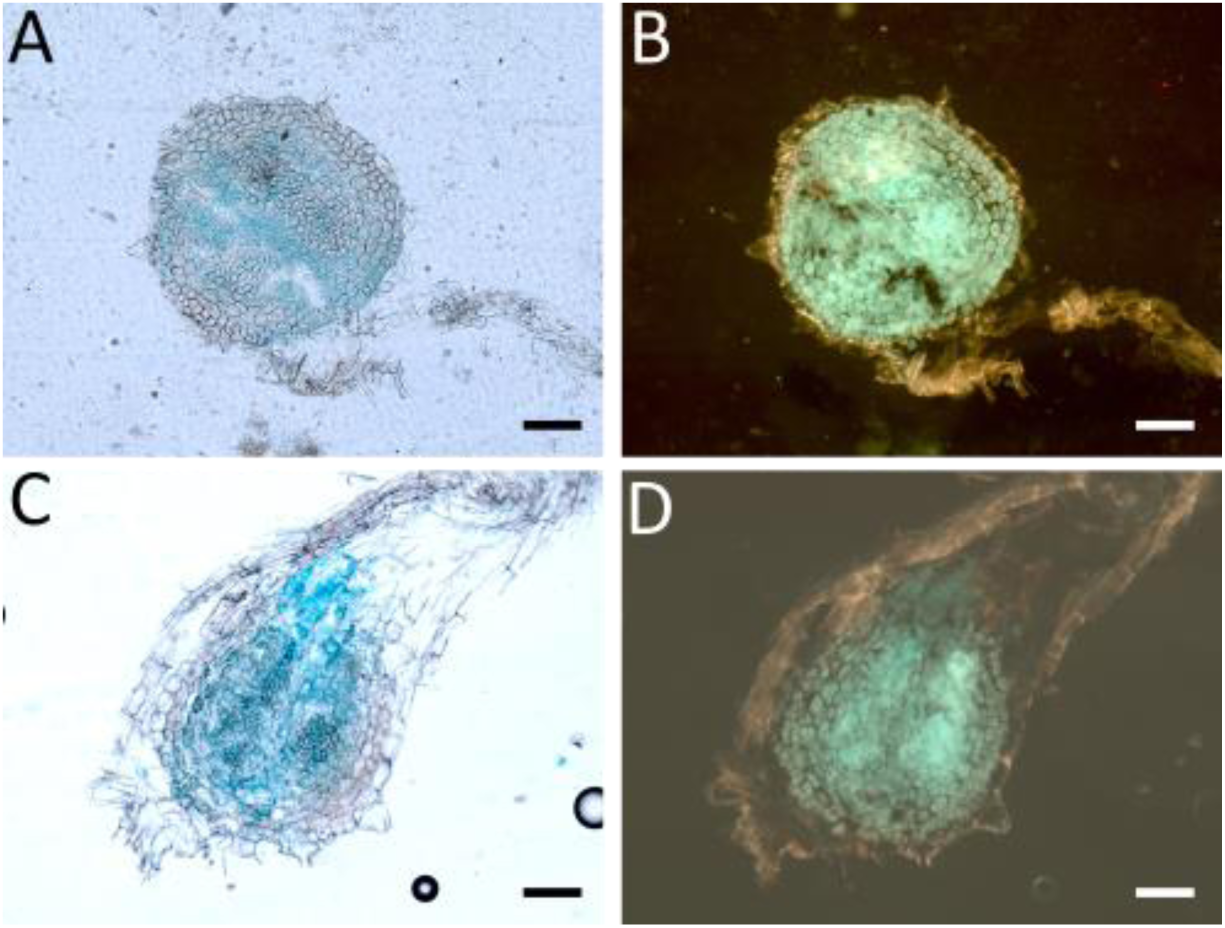
D*R*5*::GUS* staining pattern seen throughout *coch* mature nodules shown in Figure 6H. Transverse sections of *coch* mutant nodules showing DR5::GUS activity. (A–B) Top region and (C–D) basal region of the nodule. Panels (B) and (D) show GFP-tagged rhizobia. Scale bar = 100 µm.

